# High-content analysis reveals senescence of cultured canine adipose-derived mesenchymal stromal cells to be reversible with a novel immortalization approach

**DOI:** 10.1101/2020.07.27.217596

**Authors:** Ana Stojiljković, Véronique Gaschen, Franck Forterre, Ulrich Rytz, Michael H. Stoffel, Jasmin Bluteau

**Author notes:** <u>**Correspondence:**</u> Ana Stojiljković, Division of Veterinary Anatomy, Länggassstrasse 120, 3012 Bern.

## Abstract

In the last decades, the scientific community spared no effort to elucidate the therapeutic potential of mesenchymal stromal cells (MSCs). Unfortunately, *in vitro* cellular senescence occurring along with a loss of proliferative capacity is a major drawback in view of future therapeutic applications of these cells in the field of regenerative medicine. Even though insight into the mechanisms of replicative senescence in human medicine has evolved dramatically, knowledge about replicative senescence of canine MSCs is still scarce. Thus, we developed a high-content analysis workflow to simultaneously investigate three important characteristics of senescence in canine adipose-derived MSCs (cAD-MSCs): morphological changes, activation of the cell cycle arrest machinery and increased activity of the senescence-associated β-galactosidase. We took advantage of this tool to demonstrate that passaging of cAD-MSCs results in the appearance of a senescence phenotype and proliferation arrest. This was partially prevented upon immortalization of these cells using a newly designed PiggyBac™ Transposon System, which allows for the expression of the human polycomb ring finger proto-oncogene BMI1 and the human telomerase reverse transcriptase under the same promotor. Our results indicate that cAD-MSCs immortalized with this new vector maintain their proliferation capacity and differentiation potential for a longer time than untreated cAD-MSCs. This study not only offers a workflow to investigate replicative senescence in eukaryotic cells with a high-content analysis approach but also paves the way for a rapid and effective generation of immortalized MSC lines. This promotes a better understanding of these cells in view of future applications in regenerative medicine.

## 2 Introduction

In the last decades, regenerative medicine has seen dramatic developments and potential for new therapeutic approaches^1^. Much of this promise relies on stem cells, and the discovery of somatic stem cells (SSCs) in various tissues. In recent years, the scientific community spared no effort to elucidate the therapeutic potential of mesenchymal stromal cells (MSCs) for serious diseases such as orthopaedic disorders^2^, cardiovascular problems^3^, immune system ailments^4^ and cancer^5^, and several cell culture systems for human stem cells have become available.

As their name implies, MSCs originate from mesenchymal tissue. Irrespective of their occurrence in various body compartments, MSCs have two peculiar characteristics in common: the capability of self-renewal and the ability to differentiate into specific cell types^6^. Correspondingly, these multipotent cells are endowed with a great potential for many therapeutic approaches^7^.

The main obstacle for the diffusion of MSCs in routine clinical applications is the qualitative heterogeneity of the cell preparations for lack of reproducibility and standardization of the *in vitro* culture conditions. This implies an extensive characterization of the cells prior to any clinical application. This, in turn, requires a considerable number of cells, which need to be produced only to enable their characterization^8^. However, notwithstanding the capability of MSCs to self-renew, the expansion of these cells under *in vitro* culture conditions is limited. Despite enhanced research efforts to address this problem, scientists are unanimous in conceding that cellular ageing, also called senescence, has remained a major obstacle to the development of therapeutic applications based on MSCs^9^.

Senescence is a physiological process triggered by endogenous and exogenous stimuli. In principle, cells may either adapt and completely recover or undergo cell death as a response to stress-induced cell damage. Additionally, proliferating cells such as MSCs also have the capability to react to stressors by entering a state called cellular senescence^10^. This is an irreversible state of permanent cell-cycle arrest irrespective of sustained metabolic activity.

*In vivo*, this mechanism is related to ageing^11^, but it also affects cells cultured *in vitro*. The so-called replicative senescence is a phenomenon seen in cell cultures that stop proliferating even though ideal growth conditions are still met. Senescent cells not only undergo proliferation arrest but also exhibit characteristic morphological and physiological features, including increased nuclear and cytoplasmic volumes, enhanced activity of the enzyme β-galactosidase, decreased expression of the cell cycle regulator protein polycomb ring finger proto-oncogene BMI1 (BMI1) and telomere shortening^12^. The major corollary of replicative senescence in cell cultures is the impossibility of expanding MSCs beyond a limited number of passages. Altogether, *in vitro* cellular senescence going along with a loss of proliferative capacity is a major drawback in view of future therapeutic applications of MSCs in the field of regenerative medicine.

It has been shown that senescence in primary cells may be circumvented by means of immortalization techniques^13^. Some of these techniques consist of upregulating natural cellular pathways. In this respect, promising results have been shown by upregulating the human telomerase reverse transcriptase enzyme (hTERT)^14^ to counteract telomere shortening, or by upregulating the polycomb complex protein BMI1^15^, which is involved in cell cycle regulation^16^. In human adipose-derived MSCs (hAD-MSCs), the simultaneous overexpression of hTERT and BMI1 following consecutive exposure to two different lentiviral vectors has turned out to provide the best immortalization efficiency, in combination with a low impact on the cell phenotype^17^.

Even though insight into the mechanisms of senescence in human medicine has evolved dramatically^18^, knowledge about replicative senescence of canine (*Canis lupus familiaris*) MSCs is still very scarce^19–21^. Therapeutic use of stem cells in veterinary medicine has not yet progressed beyond pioneering work^22,23^. Correspondingly, there is a great interest in this field^24^, both for the importance of dogs as a model organism for human diseases in pre-clinical studies and for the relevance of dogs as a companion animal.

The aim of the present study was to provide specific knowledge on canine MSCs by characterizing canine AD-MSCs (cAD-MSCs) by means of a high-content analysis workflow. Our results show that these cells undergo phenotypical changes associated with replicative senescence similar to those that occur in their human counterparts. Furthermore, we designed a non-viral PiggyBac™ transposon system containing the human TERT and the human BMI1 coding sequences under the same promotor. This innovative approach enabled us to immortalize the cells and to prevent, to some extent, the occurrence of the senescent phenotype in cAD-MSCs.

## 3 Results

### 3.1 Proliferation rate of cAD-MSCs decreases with increasing number of passages

The proliferation assay corroborated a noticeable reduction in proliferation potential in cAD-MSCs over passages (five dog donors, age range 0.5-8 years). The number of nuclei counted at *day in vitro* 7 (DIV7) decreased progressively (Figure 1a), and cAD-MSCs reached proliferation arrest at P9. This prevented the performance of any experiments at later passages. The proliferation characteristics at different passages were compared by quantifying the area under the experimental proliferation curve (eAUC) and the carrying capacity (*K*), which both first increased between P2 and P3, but then significantly decreased at every later passage measured (Figure 1b and 1c). Consistently, the immortal control HeLa cells did not undergo proliferation arrest even at higher passages, while HeLa cells treated with 1 μM of the cytostatic drug camptothecin at DIV1 (HeLaSen) showed immediate proliferation arrest.

**Figure 1.**
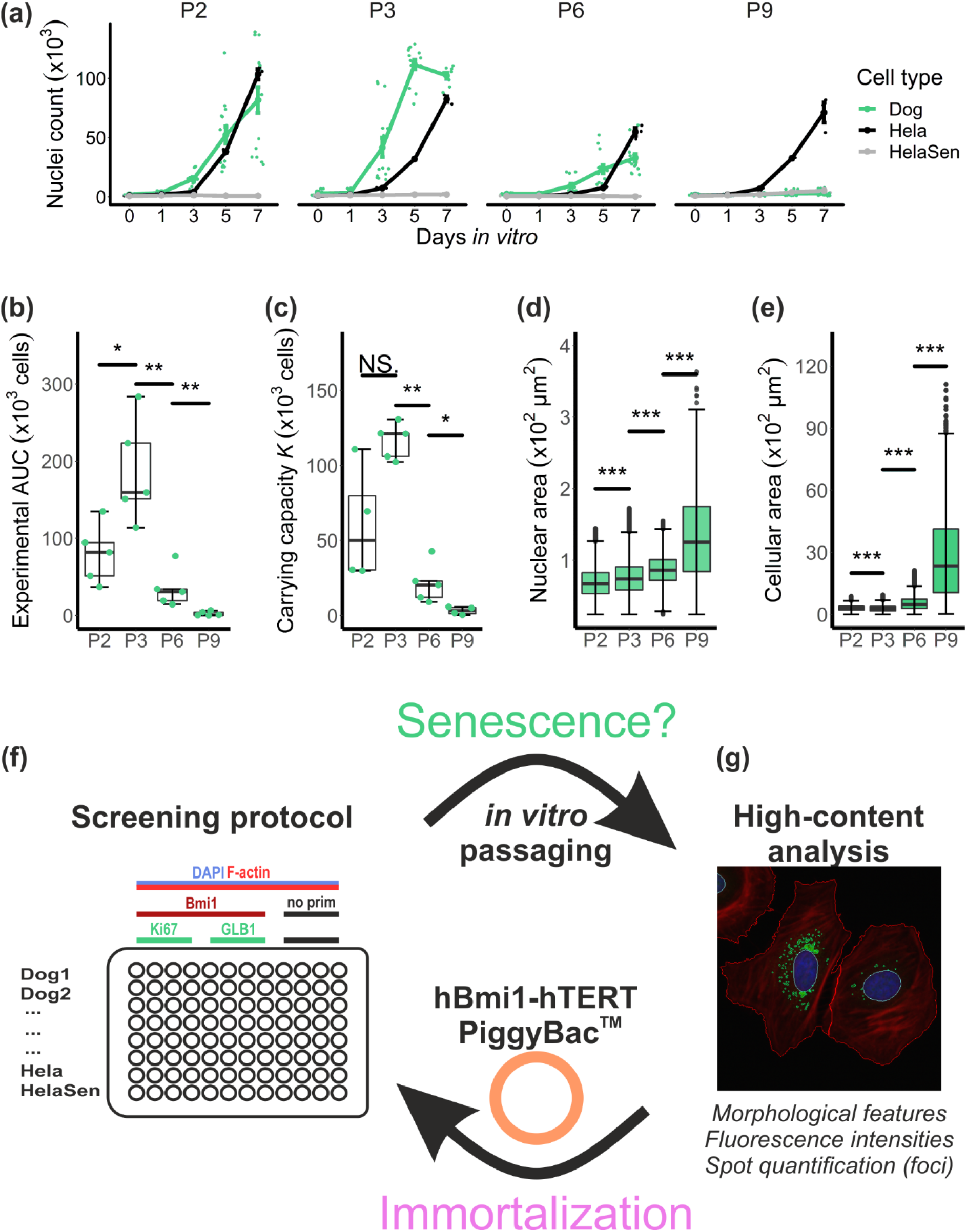
The proposed high-content analysis workflow demonstrates that *in vitro* passaging induces proliferation arrest and increased cell and nuclear size in cAD-MSCs. Logistic equations were fitted to cellular growth data (a), obtained by automated counting of nuclei in DAPI-stained micrographs over 7 days at different passages (P2-P9) (n=5 biological replicates, mean ± s.e.m.). HeLa cells and HeLa cells treated with 1 μM camptothecin to induce senescence were assessed in parallel. The progressive reduction of the extrapolated area under the experimental proliferation curve eAUC (b) and the carrying capacity *K* (c) are in agreement with a decrease in proliferation capacity starting from P3. The analysis of cell morphology features revealed a significant increase of both nuclear (d) and cellular (e) size (2D-projected area) over passages. The high-content analysis workflow consists in a four-colour immunostaining and imaging screening method to study replicative senescence in a quantitative manner (f). cAD-MSCs that were obtained from five tissue donors and two controls (HeLa and HeLaSen) were processed and imaged in the same multi-well plate. Gene expression at the protein level of three proteins involved in replicative senescence was studied by immunostaining (BMI1, Ki67 and GLB1). Incubations without primary antibodies served as controls. DAPI staining of DNA and phalloidin staining of F-actin were used to identify morphological features of cAD-MSCs (g). The acquired micrographs were automatically segmented and cellular features were extracted and quantified using open-source software exclusively. After assessing senescence in cAD-MSCs over four passages, the efficacy of immortalization using a BMI1-hTERT-PiggyBac™ transfection vector was corroborated by repeating the screening procedure on “immortalized” cAD-MSCs. NS p > 0.05; * p < 0.05; ** p < 0.01; *** p < 0.001. Scale bar = 50 μm.

### 3.2 Late-passage cAD-MSCs present characteristic morphological features of senescent cells

In this study, the potential of high-content analysis was exploited to provide a complex picture of the effects of passaging on cAD-MSCs. To this end, we developed a workflow that allowed the simultaneous analysis of cells originating from different tissue donors, three proteins involved in senescence and numerous morphological features (Figure 1f). This parallelization considerably reduced the experimental variability between samples. The workflow was used first to assess the appearance of a replicative senescence phenotype in cAD-MSCs, and later to evaluate the efficacy of a newly designed immortalization protocol to generate cAD-MSC lines (Figure 1f and 1g). The image analysis pipeline included automated nuclei and cell segmentation, and the quantification of fluorescent signals. For this purpose, only open-source software was used (segmentation pipelines available as Supplementary Table S2).

Nuclear and cellular areas measured in DAPI- and phalloidin-stained micrographs significantly increased in cAD-MSCs over each successively analysed passage. Over the entire experiment (P2-P9), a 2-fold increase in nuclear area and a 9-fold increase in cell area were observed (Figure 1d and 1e). In contrast, there were no significant differences in the measured parameters between cAD-MSCs from different tissue donors within the same passage (Supplementary Figure S1a and S1b). This allowed the pooling of cAD-MSCs results for further analysis. The enlarged nuclear and cellular areas of cAD-MSCs were already evident by visual inspection of the micrographs at different passages (Figure 2a). The same was true for camptothecin-treated control HeLaSen cells, while untreated HeLa cells did not show morphological changes upon passaging (Supplementary Figure S1c).

**Figure 2.**
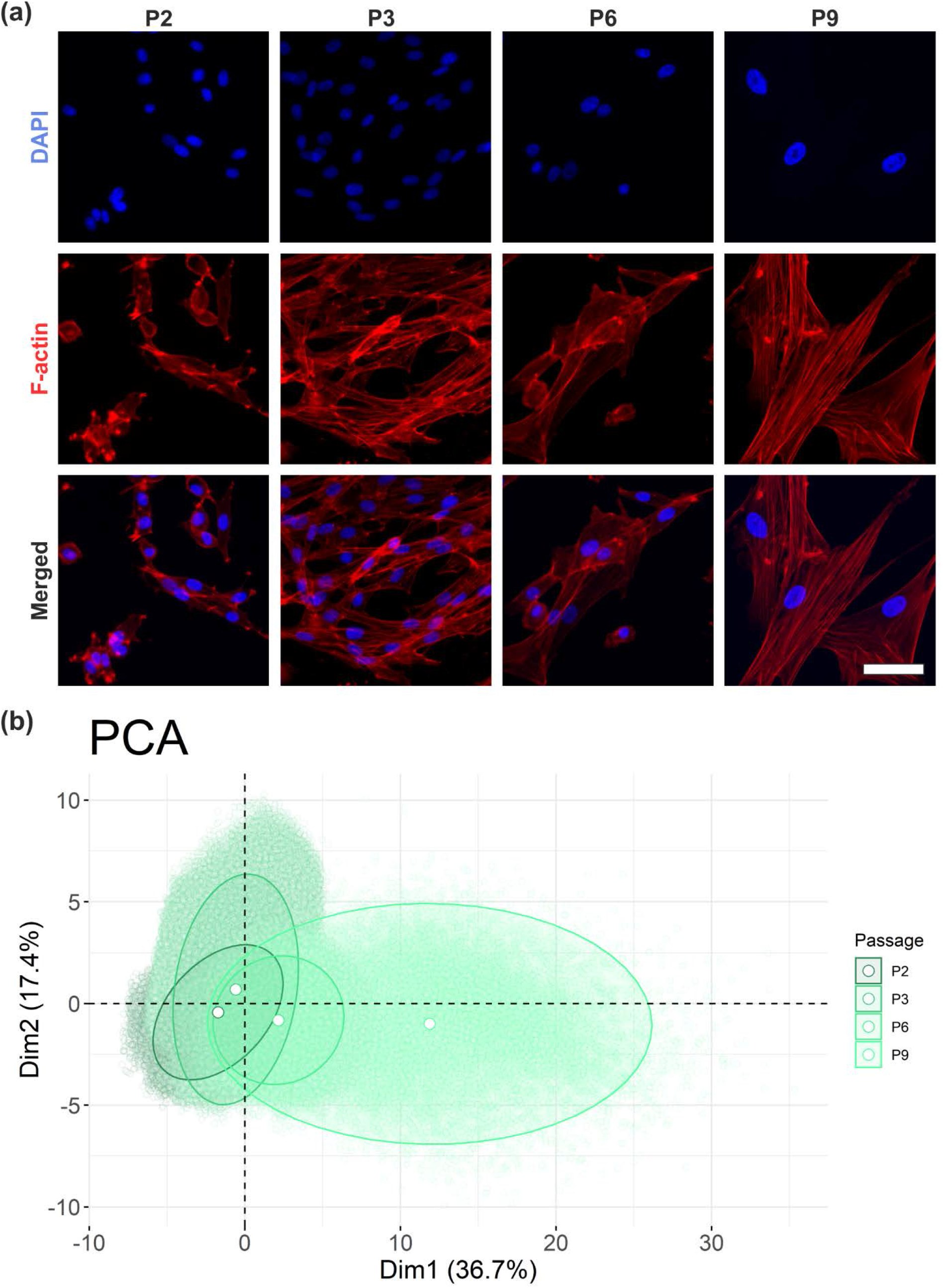
High-content analysis reveals that cellular morphological features of cAD-MSCs changed upon *in vitro* passaging. The significant increase of cell and nuclear size of cAD-MSCs is obvious by visual inspection of representative DAPI- and phalloidin-(F-actin)-stained micrographs displayed at identical magnifications (a). The dimensionality reduction (b) by principal component analysis (PCA) of the extracted morphological features revealed that cAD-MSCs assume distinct morphological characteristics during passaging, which show only minor overlap between P2 and P9. The five canine samples were pooled. The axes represent the first two dimensions of the PCA, which together explained 54.1% of the total variance. Each dot in the graph represents a single cell.

The evolution of the morphological appearance as a function of increasing numbers of passages was highlighted by principal component analysis (PCA) of defined nuclear and cellular morphological features (Supplementary Table S2). Over passages, cAD-MSCs mostly showed a right-shift on the first principal component axis (Dim1), which explained 36.7% of the variance in the experiment. Features such as cellular perimeter, cellular diameter, cellular area and nuclear area yielded the major contribution to Dim1 (Supplementary Figure S2a). The clusters of cAD-MSCs at different passages plotted in the first two dimensions showed an evident separation. In particular, cells at P9 had a small overlap with cells at earlier passages (Figure 2b). This method was used later to compare the effects of immortalization on cAD-MSCs.

### 3.3 Protein expression of cAD-MSCs at late passages was in accordance with a senescence phenotype

The high-content analysis workflow included the immunostaining of two important proteins involved in the regulation of the cell cycle, antigen Ki67 (Ki67) and BMI1. Median intensities per cells where compared, resulting in an important increase in the intensities of both proteins from the freshly thawed passage P2 to the next highly proliferative passage P3 (Figure 3b and 3d). However, we measured a significant reduction in the intensity signals for both proteins between P3 and P6. Between P6 and P9, only the BMI1 signal increased significantly. To assess the number of cells expressing the two proteins of interest, we also analysed the number of positive Ki67 and BMI1 cells by setting a threshold at the highest measured background signal (Supplementary Figure S2b and S2c). For both proteins, we observed the same trend as described for the median signal intensities, suggesting that loss of signal occurred along with loss of positive cells.

**Figure 3.**
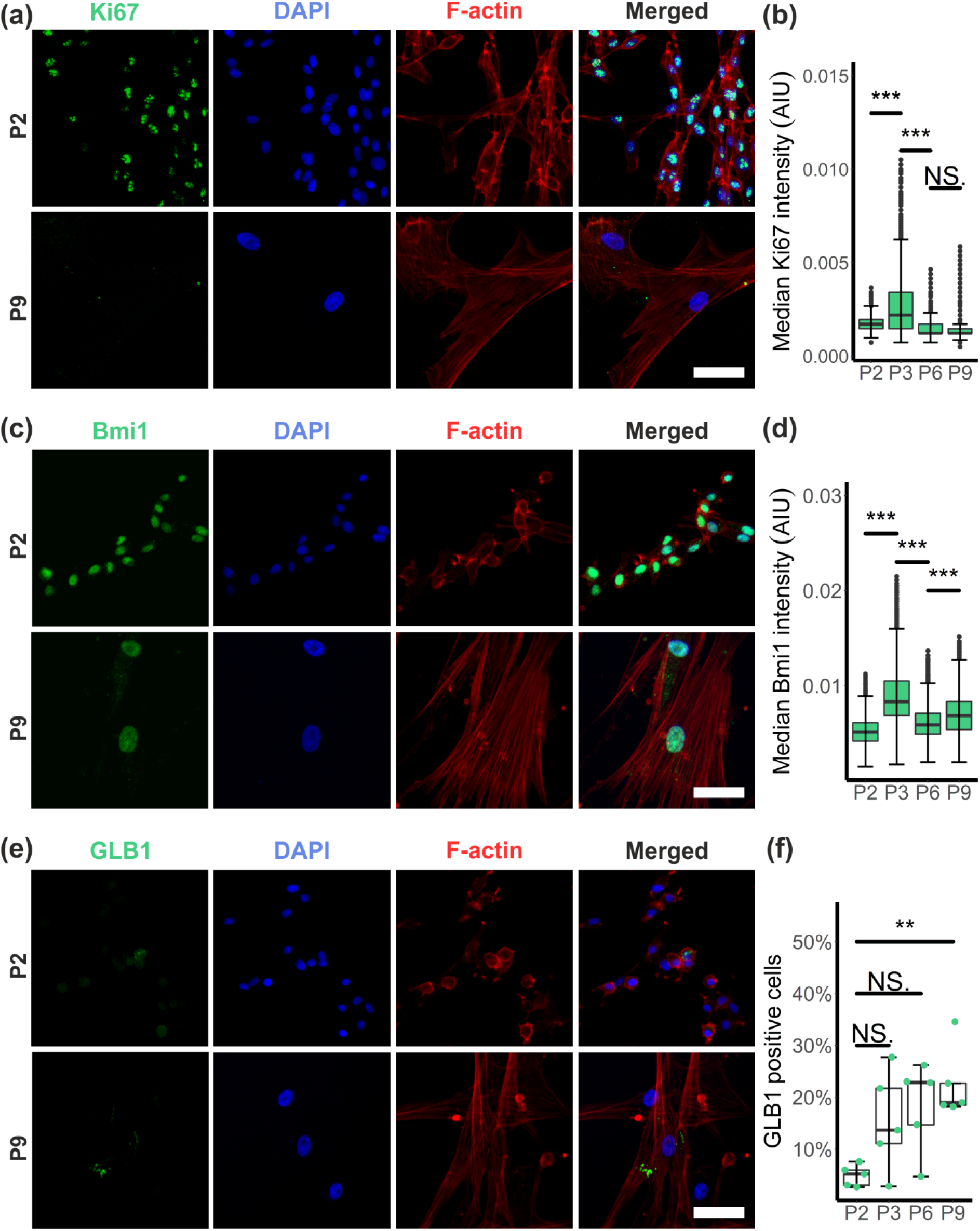
High-content analysis of the immunostaining for the detection of replicative senescence-associated proteins suggests the development of a senescent phenotype in cAD-MSCs upon *in vitro* passaging. Representative micrographs of Ki67 immunostaining at P2 and P9 (a) as well as quantification of median Ki67 intensity per cell (b) revealed an important decrease of Ki67 signal between P3 and P9. In contrast, micrographs of BMI1 immunostaining (c) suggest the maintenance of important BMI1 signal levels at P9. This visual impression was confirmed by signal quantification, which showed a fluctuation of the BMI1 signal over passages with a considerable decrease between P3 and P6 (d). GLB1 foci appear in micrographs as a vesicular accumulation of signal in the perinuclear cytoplasm (e). The number of cAD-MSCs containing at least one GLB1 focus increased progressively from P2 to P9 (f). AIU, arbitrary intensity unit. NS p > 0.05; * p < 0.05; ** p < 0.01; *** p < 0.001. Scale bars = 50 μm.

An extensively described characteristic for replicative cellular senescence is the increase in the SA-β-galactosidase enzyme activity over passages. In our experimental setup, the activity was measured indirectly by quantifying the number of cells expressing β1-galactosidase (GLB1) and the number of GLB1-signal foci per cell, as corroborated by immunostaining of this protein. Between P2 and P9, an increasing tendency toward high numbers of GLB1-positive cells was noted. Nevertheless, when comparing P9 with P2, the number of GLB1-expressing cells reached significantly higher values (Figure 3f). Within the cells presenting GLB1 foci, a 2-fold increase in cells containing a “very high” (n>20) number of foci per cell was observed at P9 (2.39%) compared to P2 (1.19%), and a moderate positive correlation between the percentage of GLB1-positive cells and the mean number of foci per cell was measured (R = 0.73, p = 0.00026) (Supplementary Figure S2d and S2e).

### 3.4 The hBMI1-hTERT PiggyBac™ transposon system provides an efficient way to transfect cAD-MSCs

To immortalize cAD-MSCs, we designed a PiggyBac™ (PB) transposon system containing the coding sequences for human BMI1 and human TERT under the same promotor (Immo-PB). The hBMI1 and hTERT coding sequences are separated by a P2A sequence, which allows the expression of both proteins to a similar amount in transfected cAD-MSCs (Figure 4a). Additionally, to assess successful transfection, a mNeonGreen (Neon) protein was introduced upstream of the hBMI1 sequence and connected to it with a P2A sequence. The transfection efficacy of a cAD-MSCs-sample at P3 with the Immo plasmid co-transfected with a hyperactive PB transposase was quantified by automated segmentation (Figure 4b). The number of Neon-expressing cells amounted to 10.7% (± 2.7%) (Figure 4c). A further advantage of this plasmid is the presence of a puromycin-resistance coding sequence, which allows for the selection of the efficiently transfected cells. After puromycin selection, the number of positive cells in the sample reached 74.5% (± 5.4%) at P4 (Figure 4c). To monitor the successful transcription of the introduced genes, we performed quantitative PCR using primers for Neon, hTERT and the transition region between hBMI1 and hTERT, as schematically represented in the plasmid map (Figure 4a). mRNA samples of Immo-transfected cAD-MSCs were collected at P6 and P10 and compared to cAD-MSCs transfected with a control PB plasmid containing only the coding sequence for the green fluorescent protein (GFP). The PCR results demonstrated that the Immo-transfected cells expressed the three tested sequences at the mRNA level, even at P10. In contrast, the cells transfected with GFP-PB only lacked the transcription of these three sequences (Figure 4d). Furthermore, we corroborated the transcription of the canine BMI1 by designing a primer that showed no binding to the human BMI1 sequence. As expected, all the mRNA samples contained the canine BMI1 mRNA. Interestingly, Immo-transfected cells allowed for the collection of enough material to perform good quality PCR for many genes even at P10, while the cells transfected with GFP-PB permitted the harvesting of only small amounts of mRNA at P10, which resulted in a very weak canine BMI1 signal and an irrelevant signal in the reference gene Glyceraldehyde 3-phosphate dehydrogenase (GAPDH) sample. The harvesting problem was confirmed with the results of the proliferation assay. In particular, at P9, cAD-MSCs transfected with GFP-PB only (GFP Dog) showed no significant increase in cell numbers over 7 days *in vitro*, while the growth curve of cAD-MSCs transfected with the Immo plasmid showed exponential growth even at P9 (Figure 4e).

**Figure 4.**
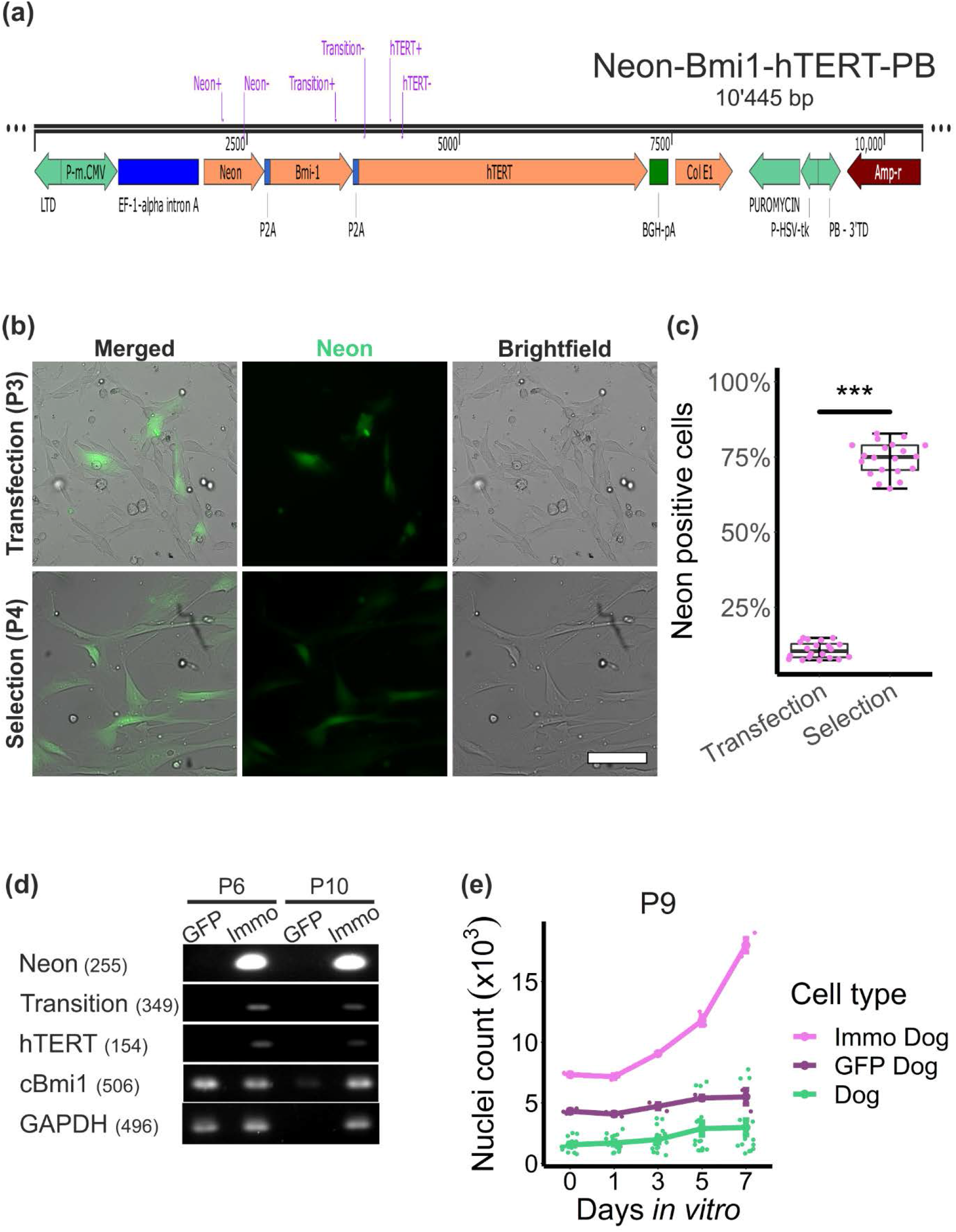
The BMI1-hTERT-PiggyBac™ transposon system successfully transfected cAD-MSCs and allowed for the puromycin selection of mNeonGreen-expressing cells. A PiggyBac™ transposon system (PB) was designed to contain the coding sequence for mNeonGreen (Neon), human BMI1 and human TERT on the same promotor (Immo-PB) (a) and was used to transfect cAD-MSCs. The Immo-PB vector consents puromycin selection of successfully transfected cells. Transfection and selection efficiency were quantified by automated counting of Neon-expressing cells and were compared between P3 at 24 h after transfection and P4 after three rounds of puromycin selection (b and c). Qualitative PCR was performed using primers for coding sequences contained in the Immo-PB (the primers are highlighted in purple in the vector map (a)). The transcription of Neon, hTERT and the transition region BMI1-hTERT were corroborated. The transcription of canine BMI1 was assessed using primers specifically designed for the canine BMI1 gene sequence, and GAPDH was assessed as a reference gene (d). cAD-MSCs transfected with the Immo-PB (Immo Dog) as well as with the control GFP-PB only containing the coding sequence for green fluorescent protein (GFP) (GFP Dog) were collected for PCR at P6 and P10. The PCR revealed that cAD-MSCs maintained the expression of the genes contained in the Immo-PB vector up to passage 10. Canine BMI1 expression was detected until later passages under all conditions. The efficacy of the Immo-PB vector to immortalize primary cells was assessed by performing the proliferation assay previously described for cAD-MSCs (e). This experiment corroborated that cAD-MSCs show important proliferation only when transfected with the Immo-PB (Immo Dog). This was not the case for untreated cAD-MSCs (Dog) or GFP-PB transfected cAD-MSCs (GFP Dog) (mean ± s.e.m.). *** p < 0.001. Scale bar = 100 μm.

### 3.5 cAD-MSCs transfected with the hBMI1-hTERT-PiggyBac™ transposon system develop a senescence phenotype, which partially recovers over passages

The high-content analysis workflow was run on Immo-transfected cAD-MSCs at P6 and P9 and compared to the results previously obtained from untreated cAD-MSCs (dog). Both nuclear and cellular areas were significantly higher in immortalized cAD-MSCs (Immo dog) than in dog cells, but in contrast to what we observed in dog cells, the immortalized cells showed a significant decrease in both parameters over passages (Figure 5a). The violin plots highlighted a skewed distribution towards smaller nuclear and cellular parameter, which was more pronounced in Immo dog cells. This is in accordance with the observation that the immortalized cells presented an inhomogeneous population in the DAPI- and phalloidin-stained samples, in which widely scattered cells showed a moderate-sized nucleus, or more often small cells with small nuclei (Figure 5b). The quantification of the Neon signal at P6 and P9 revealed that 100% of the cells expressed the fluorescence reporter gene. The PCA including the immortalized cells showed a high overlap between the immortalized cells and the P9 dog cells. The immortalized cells were even more shifted to the right on the first principal component axis, while by passaging, we observed a slight left shift instead of the right shift observed by passaging dog cells (Figure 5c).

**Figure 5.**
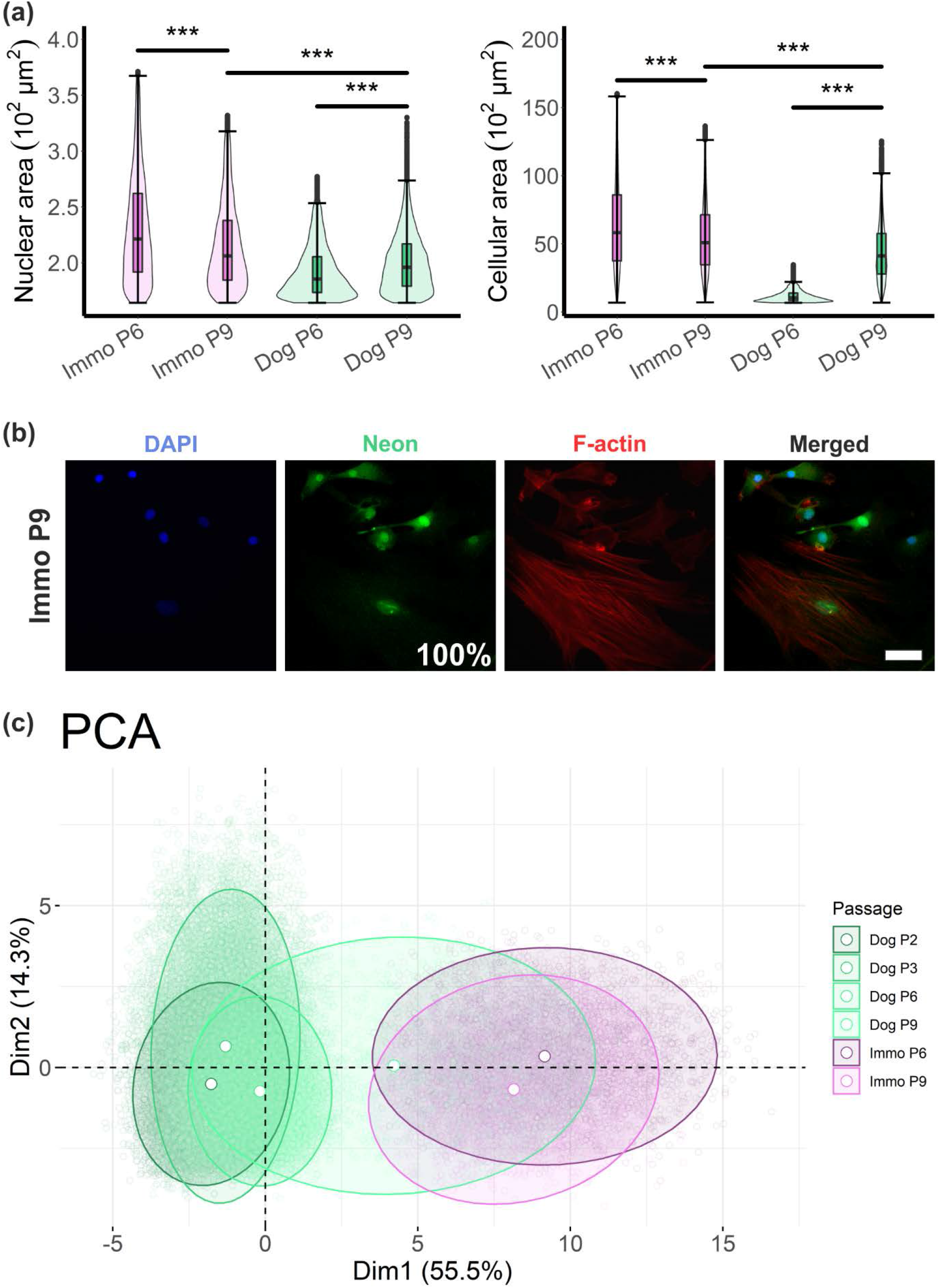
cAD-MSCs transfected with the BMI1-hTERT-PiggyBac™ show a cellular phenotype development over passages in countertendency to the untreated cAD-MSCs. When performing the high-content analysis workflow on cAD-MSCs transfected with the BMI1-hTERT-PiggyBac™ (Immo-PB) vector, we observed a significant reduction in the nuclear and cellular area between P6 and P9 (Immo cells), which is opposed to the phenotype development of untreated cAD-MSCs (Dog cells) (a). By visual inspection of the Immo cells at P9, we observed two distinct cell populations characterized by an important difference in nuclear and cellular size (b). Furthermore, the quantification of cells expressing mNeonGreen (Neon) at P6 and P9 showed an enrichment of the positive cells by passaging, without the need of further puromycin selection, reaching 100% positivity. The principal component analysis including the data of Immo cells showed that the morphological features of Immo cells at P6 and P9 are overlapping with untreated cells (Dog) mainly at P9. A slight left-shift towards the earlier passages of Dog cells is observed when comparing Immo P9 to Immo P6. NS p > 0.05; * p < 0.05; ** p < 0.01; *** p < 0.001. Scale bar = 50 μm.

Even if the cell’s morphology was comparable to that of late-passage cAD-MSCs, the expression of Ki67 and BMI1 in immortalized cells revealed a very important increase from P6 to P9 in the median intensities per cell for both proteins (Figure 6a and 6b). Furthermore, the expression of BMI1 in Immo-transfected cells at P6 was already 3-fold higher than in untreated dogs at P6, and it increased up to a factor of 10 in immortalized cells at P9. The high Ki67 and BMI1 intensities assessed with the high-content workflow were confirmed by visual inspection of the micrographs of Immo-transfected cAD-MSCs at P9 (Figure 6d and 6e). With regard to the GLB1 signal, we only have an indicative quantification of GLB1-positive cells in Immo-transfected cAD-MSCs, which was above the dog range at P6, but showed a tendency to decrease at P9, falling into the range of untreated dog cells at P9 (Figure 6c).

**Figure 6.**
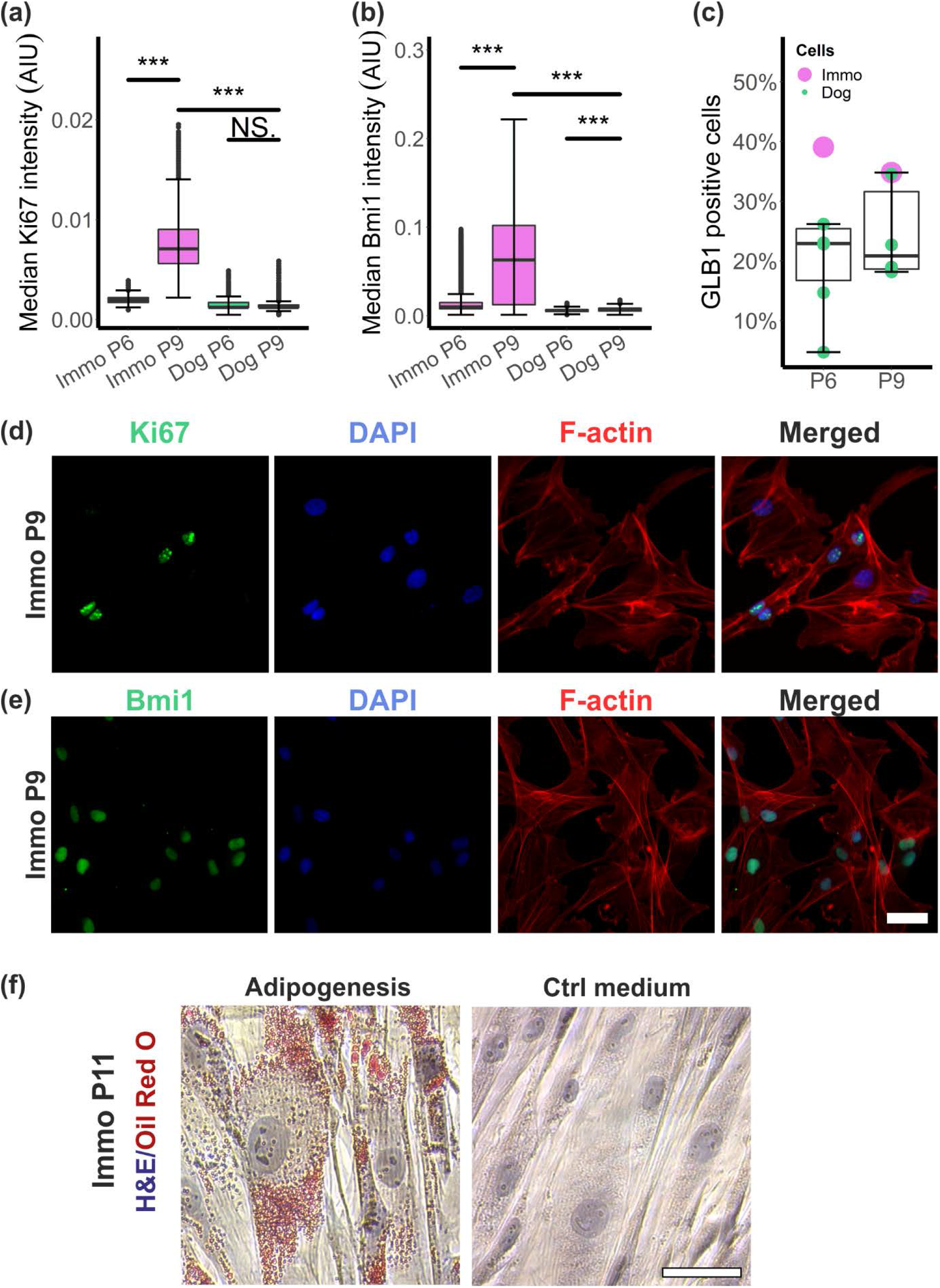
cAD-MSCs transfected with the BMI1-hTERT-PiggyBac™ show a reduction in the replicative senescence-associated gene expression over passages and maintain a high differentiation potential. In contrast to untreated cAD-MSCs (Dog), immortalized cAD-MSCs (Immo) show a significant increase in Ki67 (a) and BMI1 (b) median intensities per cell from P6 to P9. The intensity values were significantly higher than the intensities measured in Dog cells at any passage. At P9, Immo cells also show a tendency to a decreased number of GLB1-expressing cells (c). Important levels of Ki67 (d) and BMI1 (e) signals were observed by visual inspection of representative immunostaining micrographs of Immo cells at P9. Notably, MSCs at later passages lost the potential to fully differentiate into various cells of mesoderm origin (tri-lineage differentiation), i.e., adipocytes. However, Immo cells at P11 showed an important accumulation of lipid droplets (stained with Oil Red O) when exposed to adipogenesis-inducing medium (e). Control cells were cultured in standard MSC growth medium (e). AIU, arbitrary intensity unit. ns p > 0.05, *** p < 0.001. Scale bars = 50 μm.

### 3.6 cAD-MSCs transfected with the hBMI1-hTERT-PiggyBac™ transposon system maintain their differentiation potential at late passages

A main drawback of replicative cellular senescence is the loss of the differentiation potential in AD-MSCs at late passages. We, therefore, qualitatively assessed adipogenesis in Immo-transfected cells and compared it to cells transfected with GFP only. Immo-transfected cells at P11 showed a considerable accumulation of lipid droplets only when exposed to adipogenesis-inducing media (adipogenesis), as revealed with Oil Red O but not after exposure to standard growth medium (ctrl) (Figure 6f). In contrast, cells that were transfected with GFP only and which showed proliferation arrest at P9 never reached the cell density needed to start an adipogenesis experiment, and the rare cells exposed to the adipogenic medium were lost during the differentiation protocol.

## 4 Discussion

In this study, we present a high-content screening and analysis workflow relying on open-source image analysis pipelines, which was implemented to detect cellular senescence in eukaryotic cells. The workflow was first applied to investigate the appearance of replicative cellular senescence in cAD-MSCs, and later to assess the recovering from senescence of the same cell type upon treatment with a newly designed immortalization protocol.

The development of this high-content analysis workflow was based on multicolour high-throughput imaging. Taking advantage of immunostaining, quantification of the expression of three different proteins was performed simultaneously. The use of a multi-well system allowed for the reduction in the variability between experiments, as it allowed for the analysis of cAD-MSCs extracted from different tissue donors and control cells during an identical experimental run. One experimental run generated data for the quantification of numerous nuclear and cellular features and simultaneously assessed the expression of the three proteins associated with *in vitro* proliferation arrest and senescence (Ki67, BMI1 and GLB1).

The canonical SA-β-gal chromogenic assay was replaced in our study by GLB1 immunostaining. The expression of this enzyme at the protein level was reported to be strongly associated with SA-β-gal activity^25^, and, therefore, we considered its immunostaining a reasonable alternative to assessing the senescence phenotype in cAD-MSCs. The use of fluorescence-conjugated antibodies and spot-quantification instead of the classic chromogenic assay^26^ has the advantage of being less susceptible to pH fluctuations. Overall, we consider the quantification of GLB1 signal to be more suitable and reliable for high-throughput screening.

The proliferation capacity of cAD-MSCs was also quantified using high-content screening and analysis. The advantage of this system, compared to the widely accepted MTT (3-(4,5-dimethylthiazol-2-yl)-2,5-diphenyl-2H-tetrazolium bromide) proliferation assay, is the direct quantification of the cell number instead of an indirect inference from measuring the enzymatic activity by quantification of the luminescence signal, which can be affected by different extrinsic and intrinsic factors^27^. This is even more important when assessing senescence, as senescence induces important metabolic changes in the cells. Therefore, we consider the proposed high-content analysis method to be a more specific tool to assess proliferation.

The high-content analysis workflow described above was used to study replicative cellular senescence of cAD-MSCs. Proliferation arrest due to replicative senescence detected by SA-β-gal kits has been suggested previously for cAD-MSCs^19,20^, but an actual quantification of the senescence phenotype was missing for the canine species so far. Our results confirmed the occurrence of a proliferation arrest in cAD-MSCs upon *in vitro* passaging. The arrest was reached at P9, independently from the origin of the cells. Therefore, our data suggest that differences in tissue donors seem to play a minor role in the development of the senescence phenotype when compared to the effects induced by *in vitro* passaging itself. However, the indepth assessment of differences between the donors was not the main objective of this study. In general, the observed proliferation arrest at P9 is in line with previous work on human AD-MSCs^28^ and canine AD-MSCs^19^. This observation was also confirmed by measuring a significant reduction of the proliferation marker Ki67^29^ at late passages.

With the same workflow, we were able to trace the morphological evolution of cAD-MSCs over passaging and corroborated the appearance of characteristic morphological modifications as being typical of senescent cells. The main parameters were an increase in nuclear and cellular size^18,30^. We further demonstrated an important downregulation of BMI1 from P3 to P6. This protein is crucial in the maintenance of cell cycling and prevents senescence via the p16Ink4A-retinoblastoma (RB) pathway^31,32^. However, we observed an increase in BMI1 positive cells between P6 and P9, but this needs to be interpreted with caution, taking in consideration that the cells reaching P9 are probably already enriched for expressing high levels of BMI1. Furthermore, we demonstrated an increased expression of GLB1 at late passages, which provides indirect evidence for the upregulation of the β-galactosidase activity, a typical feature of senescent cells^33^.

Thus, we developed a workflow to simultaneously investigate three important characteristics of senescence^34^ and demonstrated that cAD-MSCs at late passages show all three of these characteristics: lack of proliferation markers, activation of the cell cycle arrest machinery and increased activity of the SA-β-galactosidase.

Performing this investigation on cAD-MSCs, we also observed an effect of freezing on the proliferation capacity of MSCs. When characterizing the freshly thawed cells at P2, we observed a significantly lower proliferation rate than that measured for the cells at the very next passage, P3. This observation was corroborated by both growth curve quantification and Ki67 fluorescence intensity. Furthermore, cAD-MSCs at P2 showed a reduced expression of BMI1 when compared to P3. However, the morphological features and the expression of GLB1 were only marginally affected by the freezing process. Thus, the present findings suggest a considerable impact of freezing on the proliferation capacity of MSCs, as previously described^35^, but only a minor importance in regard to inducing a senescent phenotype. Although freezing is a standard procedure in stem cell research, our observations substantiate a detrimental effect of thawing on early proliferation rate. This observation and the results suggesting a rapid onset of proliferation arrest demonstrate that the exploitable time window for any therapeutic application of these precious cells is extremely narrow.

In this study, we also described the generation of a newly designed immortalization system, which may be useful in the future to generate patient-specific AD-MSC lines. Immortalization of MSCs by overexpression of human telomerase reverse transcriptase (hTERT) has previously been reported^12^. hTERT prevents telomere shortening caused by the end replication problem, which is an important mechanism of replicative senescence^36^. Interestingly, overexpressing hTERT and BMI1 simultaneously shows more efficient immortalization of hAD-MSCs than hTERT alone^17^. A combined hTERT-BMI1 immortalization approach by using a PiggyBac™ (PB) overexpression system has been previously described for bovine fibroblasts^37^. Nevertheless, to the best of our knowledge, these two genes have always been introduced sequentially into the cells using distinct vectors. Sequential transfection has the important disadvantage of being tedious and time-consuming, as the selection of cells expressing both genes in sufficient amounts has to be warranted. Thus, this is the first report describing the use of a non-viral integrative vector to simultaneously overexpress the human BMI1 and the human TERT gene in MSCs, which might better support immortalization. In addition, this system allows for the possibility of footprint-free excision of the inserted transgene at a later point in time^38^.

Using the proposed high-content analysis workflow, we demonstrated that cAD-MSCs transfected with this new immortalization PB transposon system showed high-proliferation capacity at P9 and displayed high levels of Ki67 and BMI1 when compared to untreated cells. The expression of these two proteins increased over passages, while GLB1 expression decreased. Nevertheless, the analysis of morphological features in immortalized cAD-MSCs showed enlarged nuclear and cellular size, comparable to those of senescent cAD-MSCs cells, but these parameters decreased over passages. We hypothesize that the morphological appearance of immortalized cells briefly after immortalization is related to stress-induced senescence due to puromycin selection. In fact, it has been shown that the expression of the puromycin gene can activate pathways in the cells that lead to reactive oxygen species production^39^. Therefore, we consider the morphological appearance of immortalized cells to be mainly stress-related, and not directly related to replicative senescence, this observation being in accordance with persistently high levels of Ki67 and BMI1. Furthermore, we observed that passaging reduces the senescence phenotype observed in immortalized cAD-MSCs, probably by gradual selection of small highly proliferative cells over enlarged senescent cells. To reduce the stress-induced senescence problem, other selection techniques might be more adequate, such as flow cytometry sorting of Neon-expressing cells. Our results indicate that irrespective of their increased size, immortalized cAD-MSCs maintain differentiation potential up to late passages, as corroborated by successful adipogenesis at P11. This is very promising, as one of the main drawbacks of *in vitro* culture of MSCs is the loss of the differentiation potential over passaging, and in particular the considerable reduction in the adipogenic differentiation potential^40^.

In conclusion, we report the development of a high-content analysis workflow, which allows for the simultaneous characterization of at least three aspects of replicative senescence in eukaryotic cells in a miniaturized system. We applied the workflow to quantify the appearance of a senescence phenotype in canine adipose-derived mesenchymal stromal cells over passages and showed that this phenotype can be partially recovered by taking advantage of a newly designed non-viral immortalization vector. Many properties of these cells are still evading our understanding due to the limited possibilities of *in vitro* expansion. However, the development of more reliable immortalization techniques and, thus, the facilitated generation of patient-derived MSCs will pave the way for a better understanding of these cells and how individual variation affects their potential in view of future applications in regenerative medicine.

## 5 Experimental procedures

Unless stated otherwise, all the cell culture solutions mentioned in this chapter were purchased from ThermoFisher Scientific (Reinach, CH).

### 5.1 Cell isolation and culture conditions

Canine AD-MSCs (*Canis lupus familiaris*) were isolated as previously described for other species^41^. Briefly, five canine adipose tissue samples (Supplementary Table S1), discarded during abdominal surgeries due to indications unrelated to this study, were collected with informed owner consent in Dulbecco’s modified Eagle’s medium (DMEM) supplemented with 100 IU/ml penicillin and 100 mg/ml streptomycin (P/S) and kept at 4°C (time prior to cell isolation ranged 0.5-24 h). The sample collection was randomized by including all the adipose tissue samples discarded during surgery over one month of collection. To blind the experimenter, an identification code was assigned to the sample without any reference to the origin of the sample. The fat tissue was minced with a disposable scalpel, and the tissue fragments were enzymatically digested in 0.1% collagenase type IA (Merck, Zug, CH) in DMEM at 37°C for 1 h. After collagenase inactivation, the supernatant was filtered through a 70 μm cell strainer (Corning^®^ by Merck) and centrifuged at 500 x *g* for 10 mins. The pellet was resuspended in DMEM, and the cells were seeded in tissue-culture-treated flasks (TPP, Techno Plastic Products AG, Trasadingen, CH) in high-glucose GlutaMAX™ DMEM supplemented with 10% FBS, P/S and 10 ng/ml of human recombinant basic fibroblast growth factor (b-FGF) (PeproTech, London, UK). The cells were grown at 37°C under humidified conditions with 5% CO_2_. After 24 h, adherent cells were extensively washed with Dulbecco’s phosphate-buffered saline (DPBS), and supplied with fresh growth medium. The medium was changed every two days and passaged once until approximately 90% confluency was reached. To this end, TrypLE™ Express Enzyme was added for 5 mins. The cells were sub-cultured at 5000 cells/cm^2^.

### 5.2 PiggyBac™ transposon system plasmid preparation

A plasmid containing the hTERT sequence (#1773, Addgene, Watertown, MA, USA) was modified to contain the coding sequence for the human BMI1 followed by a P2A sequence. The human BMI1 sequence (pUC57 vector, GenScript, Leiden, NL) was inserted upstream of the hTERT sequence by enzymatic digestion (XbaI and MluI, Promega AG, Dübendorf, CH) and subsequent ligation (Rapid DNA Ligation Kit, Roche, Merck, Zug, CH). Then, we took advantage of the In-Fusion^®^ HD Cloning system (Takara Bio Europe, Saint-Germain-en-Laye, FR) to generate a PiggyBac™ transposon system (PB) containing NEON, hBMI1 and hTERT under the same promotor (Immo-PB). Briefly, we designed primers with complementary overhangs (Supplementary Table S3) and used them to generate a “NEON insert” by PCR amplification (high-fidelity DNA polymerase, CloneAmp™, Takara Bio Europe, Saint-Germain-en-Laye, FR) of the mNeonGreen sequence (Allele Biotechnology and Pharmaceuticals Inc., San Diego, CA, USA). Similarly, we generated a “hBMI1-hTERT insert” by amplifying the hBMI1-hTERT-PB plasmid previously described. Using the In-Fusion^®^ cloning system, the two inserts were introduced into a PB-backbone^42^, which was linearized by removing the EGFP coding sequence by enzymatic digestion (NotI and BsrGI, New England BioLabs, Ipswich, MA, USA) and was purified with the QIAquick Gel Extraction Kit (QIAGEN AG, Hombrechtikon, CH). DH5α competent cells were transformed with the product of the cloning reaction and the positive colonies were screened by colony PCR with a primer designed to amplify a region located between the vector backbone and the NEON insert (Supplementary Table S3).

### 5.3 Growth curves

To quantify the proliferation capacity of the cAD-MSCs, the cells were seeded in 24-well plates at a density of 5000 cells/cm^2^. For each canine sample, five plates were prepared, one for each imaging time point, including three replicate wells for each condition. At the tested time points (Day *In Vitro* (DIV) 0, 1, 3, 5, and 7), the cells were washed once with DPBS and stained with 1 μg/ml Hoechst 33342 (Merck, Zug, CH) in a Live Cell Imaging Solution (ThermoFisher Scientific) for 20 min at 37°C. After the nuclear staining, the medium was substituted with fresh medium, and 81 fields of view were imaged. Live cell imaging was performed at 37°C under humidified conditions and CO_2_-supply, taking advantage of the high-content analysis system INCell Analyzer 2000 (GE, General Electric Healthcare Europe GmbH, Glattbrugg, CH) equipped with a Plan Fluor Nikon 20x 0.45 NA objective and the 350_50x & 455_50 m (DAPI) filter set. The time point DIV0 was imaged at 4 h after the cell seeding. At this time point, the cells had enough time to adhere, but not to proliferate. The experimental setup included two types of HeLa cell controls, a genuinely immortal cell line derived from a human cervical cancer (confirmed as mycoplasma-free prior to use). HeLa cells are expected not to show signs of proliferation arrest over passages (negative control). The cells were grown in GlutaMAX™ DMEM supplemented with 10% FBS and P/S and under the same culture conditions as cAD-MSCs. The experiments were performed in parallel to the cAD-MSCs, but HeLa cells were seeded at 2500 cells/cm^2^. As a positive control for senescence, at DIV1, half of the plated HeLa cells were incubated with standard culture medium containing 1 μM camptothecin (CPT) for 3 h. After this period, the medium was changed back to the standard medium. Effects of impaired growth were expected from DIV3 on, as the plant-derived cytostatic CPT has been reported to induce proliferation arrest and senescence phenotype also in immortal cells over a mechanism of topoisomerase I-mediated DNA damage^43^.

The images were segmented using an Otsu two-class thresholding strategy with the CellProfiler™ software (version 3.8.1). For the detailed analysis pipeline, see Supplementary Table S2.

Prior to pooling the five dog samples, we verified the absence of significant differences between donors by using the Kruskal-Wallis test, followed by a Dunn-Bonferroni post hoc test (p < 0.05). A logistic regression model was fitted to the data of each dog sample using the Growthcurver package for R, and the area under the experimental proliferation curve (eAUC) and the carrying capacity *K* were extrapolated and plotted over the four passages.

### 5.4 Immunostaining and high-content analysis

For the characterization of the senescence phenotype in cAD-MSCs by means of immunostaining, the cells were seeded in Greiner μClear^®^ 96-well plates (Greiner Bio-One, Kremsmünster, AT) at a density of 5000 cells/cm^2^ (HeLa: 2500 cells/cm^2^). The cells were grown under standard culture conditions for 48 h prior to fixation and immunostaining (detailed protocol in Supplementary Table S3). For the quantification of the Ki67 and GLB1 signals, each plate contained four well replicates, while eight replicate wells were plated for the BMI1 signal.

Images were acquired with the high-content analysis system INCell Analyzer 2000 (GE, General Electric Healthcare Europe GmbH, Glattbrugg, CH) equipped with a Plan Fluor Nikon 20x 0.45 NA objective (filter sets in Supplementary Table S3). For each well, 30 randomly distributed fields of view were imaged.

The images were segmented using the CellProfiler™. For the nuclear segmentation, a similar strategy was used as previously described for the quantification of the proliferation capacity. Furthermore, the cell contours were segmented by using the nucleus as a reference object, which was propagated in the micrographs of the Alexa Fluor™ 555 Phalloidin signal (pipeline in Supplementary Table S2). Nuclear and cellular areas were compared over passages considering the tissue donors as biological replicates. Stratified randomized sub-sampling of the data was used to address the problem of p-hacking in big data^44^. We observed that sub-sampling eliminated the differences between donors, while the significant differences between passages remained unchanged. For nuclear and cellular areas, only non-parametric tests were applied because the distribution of the data showed positive skewness.

To address the question of the morphological alteration of cAD-MSCs over passages, we performed principal component analysis (PCA). For this purpose, 10 nuclear and 10 cellular features (Supplementary Table S2) were extracted from the segmented objects and analysed using the PCA function of the R package FactoMineR and the factoextra R package.

For the quantification of the immunofluorescence signal, the nuclear segmentation was used as reference object to measure the median intensity of Ki67 and BMI1 after the subtraction of the background, measured as minimal intensity value of the Ki67 or BMI1 signals, respectively. The subtraction of the background was performed to minimize the plate-to-plate effect. In this case, normalization based on DAPI was considered inappropriate due to the observed effects on nuclear morphology at different passages. For the count of GLB1-positive cells, GLB1 foci were segmented and assigned by 100% overlapping to objects resulting from cellular segmentation. To evaluate the possible accumulation of GLB1 foci over passages, the number of GLB1 foci was assigned to four categories: GLB1 foci ≤ 3 as “low”, GLB1 foci > 3 and ≤ 10 as “medium”, GLB1 foci > 10 and ≤ 20 as “high”, and GLB1 > 20 as “very high”.

### 5.5 Transfection of cAD-MSCs with the PiggyBac™ transposon system

cAD-MSCs at P2 were seeded in 24-well plates at a density of 5000 cells/cm^2^ and grown under standard culture conditions for 24 h. Then, they were transfected with 0.5 μg DNA, which was composed of a 5:1 ratio of Immo-PB and a PiggyBac™ transposase^42^. The transfection was performed with the Lipofectamine Stem reagent following the manufacturer’s instructions. At 24 h after transfection, brightfield and fluorescent images were acquired to assess the number of NEON-expressing cells. For this purpose, live cell imaging was performed with the INCell Analyzer 2000 using the environmental control, as previously described. The images were automatically segmented using CellProfiler™ (pipeline in Supplementary Table S2). After imaging, the cells were split with standard protocols and transferred to 6-well plates to allow proliferation of the successfully transfected cells. The cells were monitored for confluency, and as soon as NEON-positive colonies appeared, the cells were treated with 10 μg/mL puromycin. The medium was changed daily to remove the cell debris and replaced with fresh medium containing puromycin. After three days of puromycin treatment, the medium was replaced with standard culture medium. The medium was replaced every second day, and the cells were monitored daily. The cells required 10 days to repopulate the well and reach confluency. At this moment, they were split at a 1:3 ratio and treated for three days with puromycin, as previously described. After the last puromycin treatment, the cells were imaged to assess the number of NEON-positive cells, which reached 100% at this point. The immortalized cells underwent the exact same experimental procedures as described for cAD-MSCs. In parallel to the Immo-PB transfection, cells transfected with a GFP-only control PB were generated and used as a control. The senescence screening protocol was adapted to the fact that immortalized cells express NEON fluorescence. To this end, the number of replicates was reduced to three for the stained proteins Ki67, BMI1, and GLB1, and only single-staining was performed instead of double-staining.

Furthermore, we used standard PCR protocols to assess the expression of the inserted constructs at different passages (detailed protocol in Supplementary Table S3).

### 5.6 Adipogenesis in immortalized cAD-MSCs

The adipogenesis was performed as previously described for cAD-MSCs^45^. At P11, cAD-MSCs transfected with either Immo-PB or the GFP-only PB control were seeded at 10000 cells/cm^2^ and cultured under standard conditions. After 24 h, the medium was changed to adipogenesis-inducing medium containing high-glucose DMEM (4.5 g/L glucose), 10% FBS, 5% rabbit serum, 1 μM dexamethasone (Merck), 5 μg/mL insulin (Merck) and 5 μM rosiglitazone (Merck). After 21 days of exposure to the adipogenic medium, which was replaced every three days, the cells were fixed with 2% paraformaldehyde for 20 mins. Oil Red O staining was performed as described by the manufacturer (Lonza AG, Basel, Switzerland) and was followed by haematoxylin and eosin (H&E) counterstain.

### 5.7 Data analysis

Data processing and analysis were performed with KNIME 3.7.0 (Zürich, CH) and the R environment (version 3.6.1) using R Studio version 1.2.5033 (Boston, MA, USA). Statistical analysis was performed with R, and the figures were prepared with the R package ggplot2 and assembled with CorelDRAW Graphics Suite 2018 (Corel Corporation, Ottawa, CAN). The images were prepared using FIJI (ImageJ 1.52p). Beside the graphical representation, all the relevant results are presented as numerical values in Supplementary Table S4. The image analysis pipelines are available in the following GitHub repository: https://github.com/StojiljkovicVetAna/HCA-to-investigate-senescence.

## 6 Acknowledgements

We gratefully acknowledge the kind support of the staff from the Small Animal Clinic, Vetsuisse Faculty Bern and the assistance of Helga Mogel in the lab. We also thank Philippe Plattet and Marianne Wyss for providing us with plasmids and helping with cloning. We thank Meike Mevissen, Angélique Ducray, Volker Enzmann and Simone Forterre for helpful discussions. This study was performed with the support of the interfaculty Microscopy Imaging Center (MIC) of the University of Bern.

## 7 Conflict of interest statement

The authors declare no conflict of interest.

## 8 Author contributions

AS designed the high-content analysis approach, wrote the manuscript and created the figures with the support of JB and MHS. AS performed all the experiments with the help of VG for cell culture and molecular biology. FF and UR organized the collection of the tissue samples.

## 9 Data availability statement

The data that support the findings of this study are available from the corresponding author upon reasonable request.

## 11 Supporting information

- Supplementary Table S1
- Supplementary Table S2
- Supplementary Table S3
- Supplementary Table S4
- Supplementary Figure S1
- Supplementary Figure S2

## Notes

### Competing Interest Statement

The authors have declared no competing interest.

