## Supplementary Figure S1 for "High-content analysis reveals senescence of cultured canine adipose-derived mesenchymal stromal cells to be reversible with a novel immortalization approach"

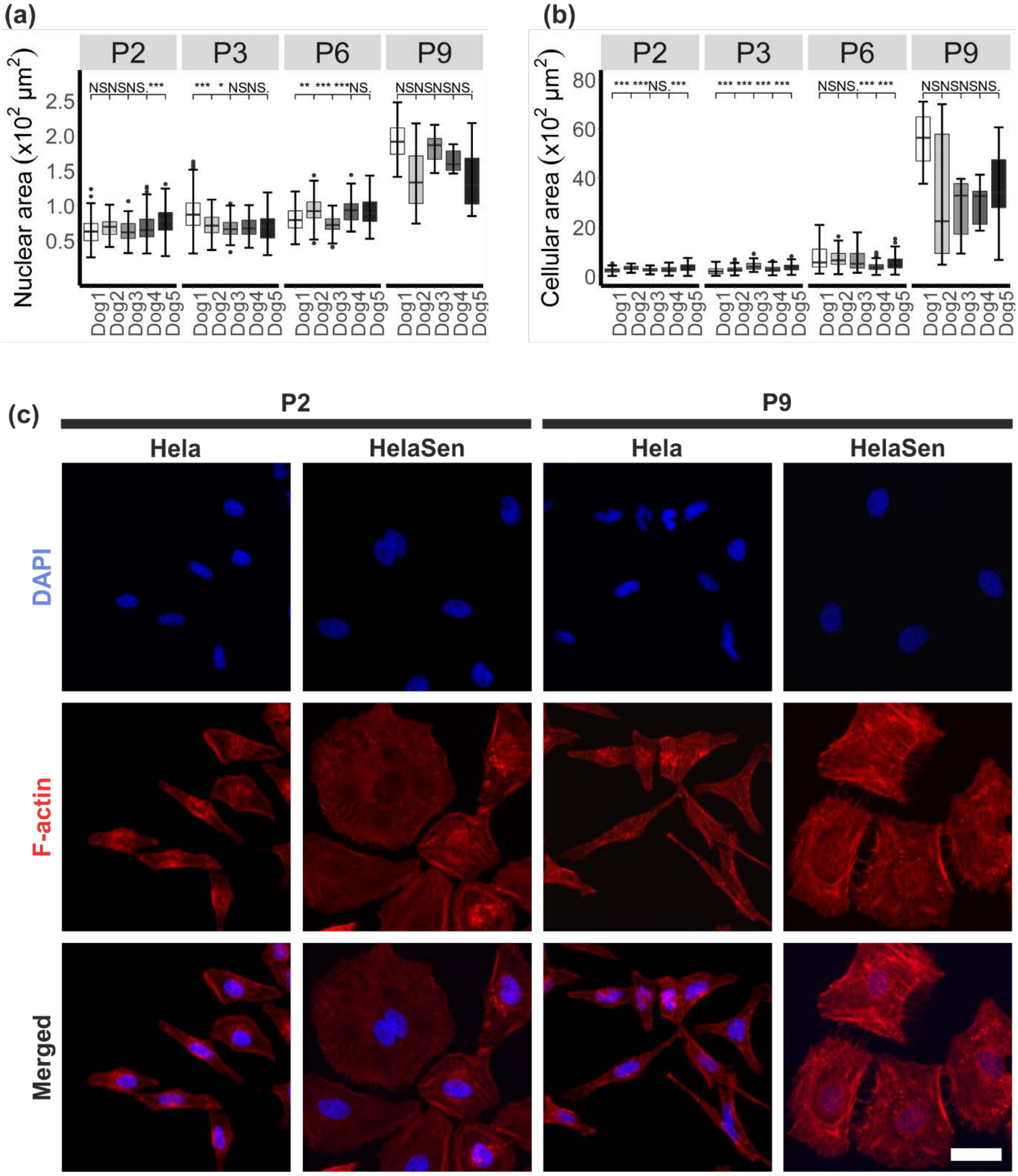
(a)
P2
P3
P6
P9
NSNSNS.***
*** * NSNS.
** *** ***NS.
NSNSNSNS.
Nuclear area (x10² μm²)
2.5
2.0
1.5
1.0
0.5
Dog1
Dog2
Dog3
Dog4
Dog5
(b)
P2
P3
P6
P9
*** ***NS.***
*** *** *** ***
NSNS.*** ***
NSNSNSNS.
Cellular area (x10² μm²)
80
60
40
20
0
Dog1
Dog2
Dog3
Dog4
Dog5
(c)
P2
P9
Hela
HelaSen
Hela
HelaSen
DAPI
F-actin
Merged

**Supplementary Figure S1**

High-content analysis of morphological characteristic predominantly revealed non-significant (NS) differences in both nuclear (a) and cellular (b) size (2D-projected area) between cAD-MSCs isolated from different tissue donors. Furthermore, when significant, the difference was usually not conserved over different passages, suggesting that different tissue donors may be considered as biological replicates. (c) The significant increase in nuclear and cellular size of Hela cells upon treatment with 1 µm camptothecin (HelaSen) is evident by visual inspection of representative DAPI- and phalloidin-(F-actin)-stained micrographs displayed at identical magnifications. Untreated Hela cells (Hela) maintain a constant morphological appearance at different passages. Therefore, we considered this cell to be an adequate negative control for replicative senescence. In contrast, HelaSen display an increased size at all analysed time points, justifying their use as positive control for the appearance of the replicative senescence phenotype. Scale bar = 50 µm. ns $p > 0.05$; * $p < 0.05$; ** $p < 0.01$; *** $p < 0.001$.
