## Supplementary Figure S2 for "High-content analysis reveals senescence of cultured canine adipose-derived mesenchymal stromal cells to be reversible with a novel immortalization approach"

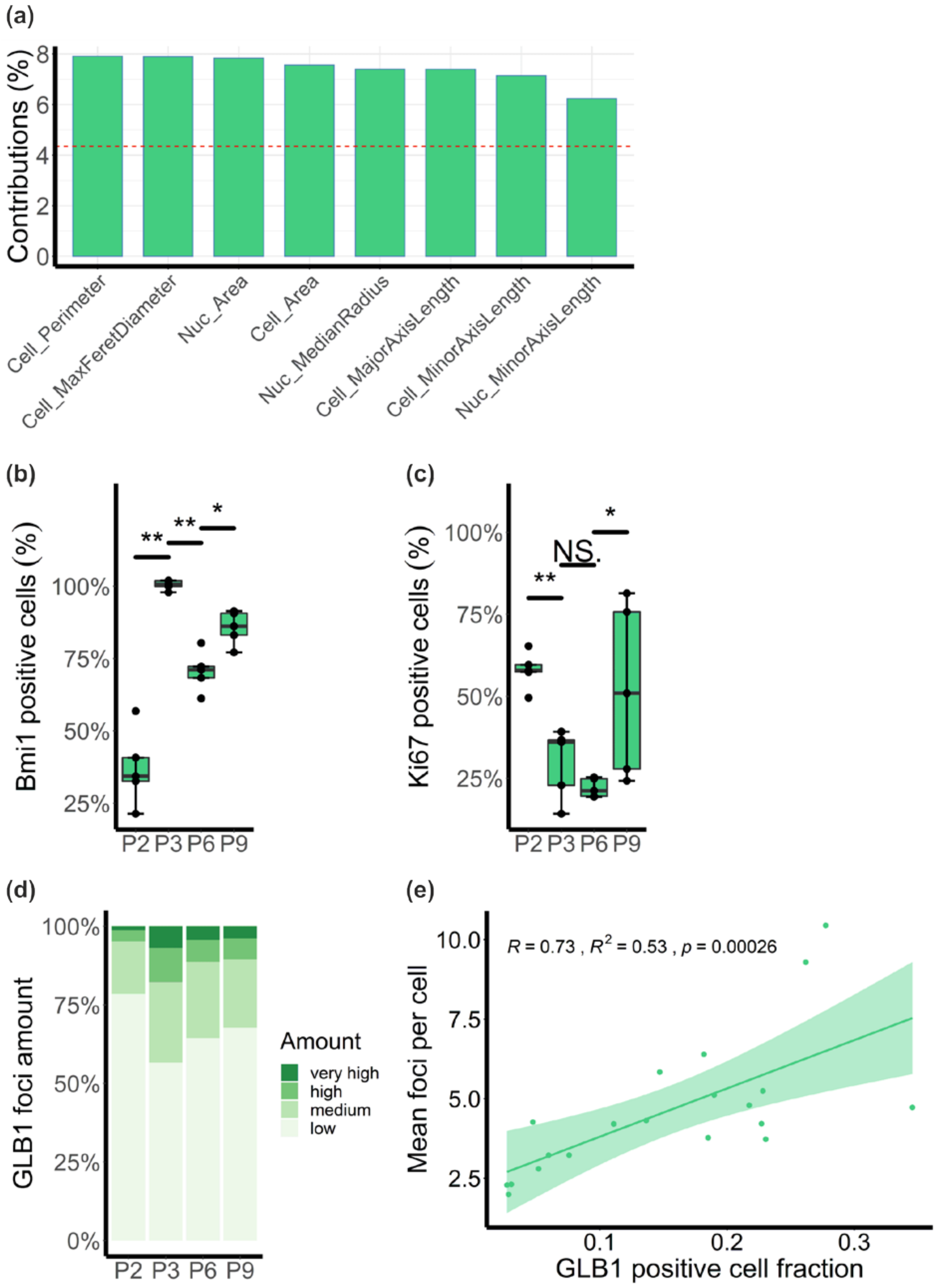
(a)
Contributions (%)
8
6
4
2
0
Cell_Perimeter
Cell_MaxFeretDiameter
Nuc_Area
Cell_Area
Nuc_MedianRadius
Cell_MajorAxisLength
Cell_MinorAxisLength
Nuc_MinorAxisLength
(b)
Bmi1 positive cells (%)
100%
75%
50%
25%
**
**
*
P2 P3 P6 P9
(c)
Ki67 positive cells (%)
100%
75%
50%
25%
**
NS.
*
P2 P3 P6 P9
(d)
GLB1 foci amount
100%
75%
50%
25%
0%
P2 P3 P6 P9
Amount
very high
high
medium
low
(e)
R = 0.73 , R2 = 0.53 , p = 0.00026
Mean foci per cell
10.0
7.5
5.0
2.5
0.1
0.2
0.3
GLB1 positive cell fraction

**Supplementary Figure S2**

(a) Contribution of the first eight variables (nuclear and cellular morphological features) to the first dimension of the principal component analysis (Dim-1). The red dashed line corresponds to the expected value if the contributions were uniform (*factorextra* R package). (b) The fraction of Bmi1-positive cells at different passages follows an identical trend as the median Bmi1 intensity per cells. (c) The same was true for Ki67. This suggests that the observed values are not only due to a reduced expression within a cell, but is rather due to decrease in the number of Bmi1- or Ki67-expressing cells respectively. (d) To better understand the occurrence of GLB1 foci in cAD-MSCs at different passages, as corroborated by immunostaining, we classified the amount of foci in the cell: GLB1 foci $\leq 3$ as "low", GLB1 foci $> 3$ and $\leq 10$ as "medium", GLB1 foci $> 10$ and $\leq 20$ as "high", and GLB1 $> 20$ as "very high". Compared to the starting point P2, we observed a general increase of cells expressing a high to very high amount of GLB1 foci at higher passages, with the major increase between P2 and P3. (e) To corroborate the relationship between the number of cells expressing GLB1 and the number of GLB1 foci, we performed a Spearman correlation analysis and found a moderately positive correlation ($R=0.73$) between the two variables. ns $p > 0.05$; * $p < 0.05$; ** $p < 0.01$; *** $p < 0.001$.
