## Supplementary Table S1 for "High-content analysis reveals senescence of cultured canine adipose-derived mesenchymal stromal cells to be reversible with a novel immortalization approach"

### Supplementary Table S1: Dog tissue samples

| *Sample ID* | *Sex*[1] | *Age(y)* | *Breed* | *Sample mass (g)* | *Surgery indication* |
|---|---|---|---|---|---|
| ***Dog 1*** | fn | 3.4 | Bullterrier | 81.7 | foreign body |
| ***Dog 2*** | f | 0.5 | Bruno Jura Hound | 14.5 | portosystemic shunt |
| ***Dog 3*** | f | 1.0 | Saluki | 7.9 | mamma tumour |
| ***Dog 4*** | f | 1.1 | Rhodesian Ridgeback | 21 | intestinal invagination |
| ***Dog 5*** | f | 8.3 | German Shepard | 44 | mamma tumour |
| ***Immo Dog*** | m | 5.4 | Flat Coated Retriever | 95 | pancreatitis |

[1] f=female, m=male, n=neutered
