## Supplementary Table S2 for "High-content analysis reveals senescence of cultured canine adipose-derived mesenchymal stromal cells to be reversible with a novel immortalization approach"

**Supplementary Table S2: CellProfiler™ Pipelines**
The following pipelines were generated using the 3.8.1 version of CellProfiler™.
The image analysis pipelines are available in the following GitHub repository:
https://github.com/StojiljkovicVetAna/HCA-to-investigate-senescence.

### Proliferation of cAD-MSCs

| Module | Parameter | Setting |
|---|---|---|
| **Metadata** | *Extract Metadata?* | Yes |
| | *Metadata extraction method* | Extract from file/folder names |
| | *Metadata source* | File name |
| | *Regular expression to extract from* | (?P<WellRow>[A-H]) - (?P<WellCol>[0-9]{2})\(fld (?P<Field>[0-9]{3}) wv (?P<Channel>[A-Z]) |
| | *Extract Metadata from* | All images |
| **NamesAndTypes** | *Select the rule criteria* | File Does Contain "DAPI" |
| | *Name to assign these images* | DAPI |
| | *Select the image type* | Grayscale image |
| | *Select intensity range from* | Image metadata |
| **Groups** | *Want to group your images?* | Yes |
| | *Metadata category* | Well |
| **CorrectIlluminationCalculate** | *Select the input image* | DAPI |
| | *Name the output image* | IllumDNA |
| | *Select how the illuiation fucntion is calculated* | Background |
| | *Block size* | 60 |
| | *Rescale the illumination function* | Yes |
| | *Calculate function for each image individually?* | Each |
| | *Smoothing method* | Median Filter |
| | *Method to calculate smoothing filter size* | Manually |
| | *Smoothing filter size* | 10 |
| **CorrectIlluminationApply** | *Select the input image* | DAPI |
| | *Name the output image* | CorrDNA |
| | *Select the illumination funciton* | IllumDNA |
| | *Select how the function is applied* | Divide |
| **IdentifyPrimaryObjects** | *Use advanced settings* | Yes |
| | *Select the input image* | CorrDNA |
| | *Primary objects to be identified* | Nuc |
| | *Typical diameter of objects (Min; Max)* | 15;60[1] |
| | *Discard objects outside the diameter range?* | Yes |
| | *Discard objects touching the border of the image?* | Yes |
| | *Threshold strategy* | Global |
| | *Thresholding method* | Otsu |
| | *Two-class or three-class thresholding* | Two classes |
| | *Threshold smoothing scale* | 1.3488 |
| | *Threshold correction factor* | 1 |
| | *Lower and upper bounds on threshold* | 0.03;1.0 |
| | *Method to distinguish clumped objects* | Shape |
| | *Method to draw dividing lines beween clumped objects* | Propagate |
| | *Automatically calculate size of smoothing filter for declumping* | Yes |
| | *Automatically calculate minimum allowed distance between local maxima* | Yes |
| | *Speed up by using lower-resolution image to find local maxima* | Yes |
| | *Fill holes in identified objects?* | After both thresholding and declumping |

[1] This are the settings for cAD-MSCs and Hela cells. For the control HelaSen the settings are: 30;300

| | | |
|---|---|---|
| **FilterObjects** | *Select the objects to filter* | Nuc |
| | *Name the output objects* | FilterNuc |
| | *Select the filtering mode* | Measurements |
| | *Select the filtering method* | Limits |
| | *Select the measurement to filter by Category* | Area Shape |
| | *Select the measurement to filter by Measurement* | FormFactor |
| | *Minimum measurement value* | 0.3 |
| | *Maximum measurement value* | 1.0 |
| **MeasureObjectSizeShape** | *Select objects to measure* | FilterNuc |
| **RescaleIntensity** | *Select the input image* | DAPI |
| | *Name the output image* | ResDNA |
| | *Rescaling method* | Stretch each image to use the full intensity range |
| **OverlayOutlines** | *Select image on which to display outlines* | ResDNA |
| | *Name the output image* | DNAOver |
| | *Outline display mode* | Color |
| | *How to outline* | Outer |
| | *Select objects to display* | FilterNuc |
| **SaveImages** | *Select the type of image to save* | Image |
| | *Select the image to save* | DNAOver |
| | *Select method for constructing file names* | From image filename |
| | *Saved file format* | Tiff |
| | *Image bit depth* | 8-bit integer |
| | *When to save* | Every cycle |
| **ExportToSpreadsheet** | *Select the column delimiter* | Comma(",") |
| | *Add image metadata to your object data file?* | Yes |
| | *Press button to select measurements* | FilterNuc<br>- Number/Object |

### High-content analysis of morphological features and Ki67/Bmi1 expression

| Module | Parameter | Setting |
|---|---|---|
| **Metadata** | *Refer to Proliferation of cAD-MSCs pipeline -* **Metadata** | |
| **NamesAndTypes** | *Select the rule criteria* | File Does Contain "DAPI" |
| | *Name to assign these images* | DAPI |
| | *Select the rule criteria* | File Does Contain "FITC" |
| | *Name to assign these images* | Ki67 |
| | *Select the rule criteria* | File Does Contain "Cy3" |
| | *Name to assign these images* | Skel |
| | *Select the rule criteria* | File Does Contain "Cy5" |
| | *Name to assign these images* | Bmi1 |
| | *Select the image type* | Grayscale image |
| | *Select intensity range from* | Image metadata |
| **Groups** | *Refer to Proliferation of cAD-MSCs pipeline -* **Groups** | |
| **IdentifyPrimaryObjects** | *Use advanced settings* | Yes |
| | *Select the input image* | DAPI |
| | *Primary objects to be identified* | Nuc |
| | *Typical diameter of objects (Min; Max)* | 15;60[2] |
| | *Discard objects outside the diameter range?* | Yes |
| | *Discard objects touching the border of the image?* | Yes |
| | *Threshold strategy* | Global |
| | *Thresholding method* | Otsu |
| | *Two-class or three-class thresholding* | Two classes |
| | *Threshold smoothing scale* | 1.3488 |
| | *Threshold correction factor* | 1 |
| | *Lower and upper bounds on threshold* | 0.023;1.0 |
| | *Method to distinguish clumped objects* | Shape |
| | *Method to draw dividing lines beween clumped objects* | Propagate |
| | *Automatically calculate size of smoothing filter for declumping* | Yes |
| | *Automatically calculate minimum allowed distance between local maxima* | Yes |
| | *Speed up by using lower-resolution image to find local maxima* | Yes |
| | *Fill holes in identified objects?* | After both thresholding and declumping |
| **IdentifySecondaryObjects** | *Select the input image* | Skel |
| | *Select the input objects* | Nuc |
| | *Name the objects to be identified* | Cell |
| | *Select the method to identify the secondary objects* | Propagation |
| | *Threshold strategy* | Global |
| | *Thresholding method* | Minimum cross entropy |
| | *Threshold smoothing scale* | 0.5 |
| | *Threshold correction factor* | 1.0 |
| | *Lower and upper bounds on threshold* | 0.027;1.0 |
| | *Regularization factor* | 0.05 |
| | *Fill holes in identified objects?* | Yes |
| | *Discard secondary objects touching the border of the image* | No |
| **MeasureObjectSizeShape** | *Select objects to measure* | Nuc, Cell |
| **MeasureObjectIntensity** | *Select an image to measure* | DAPI, Ki67, Bmi1 |
| | *Select objects to measure* | Nuc, Cell |
| **RescaleIntensity** | *Select the input image* | DAPI |
| | *Name the output image* | ResDNA |
| | *Rescaling method* | Stretch each image to use the full intensity range |
| **RescaleIntensity** | *Select the input image* | Skel |

[2] This are the settings for cAD-MSCs. For Hela and HelaSen the settings are: 30;120.

| | | |
|---|---|---|
| | *Name the output image* | ResSkel |
| | *Rescaling method* | Stretch each image to use the full intensity range |
| **GrayToColor** | *Select a color scheme* | RGB |
| | *Select the image to be colored red* | ResSkel |
| | *Select the image to be colored blue* | ResDAPI |
| | *Name of the output image* | Fused |
| **OverlayOutlines** | *Select image on which to display outlines* | Fused |
| | *Name the output image* | SkelOver |
| | *Outline display mode* | Color |
| | *How to outline* | Outer |
| | *Select objects to display* | Nuc, Cell |
| **SaveImages** | *Select the type of image to save* | Image |
| | *Select the image to save* | SkelOver |
| | *Select method for constructing file names* | From image filename |
| | *Saved file format* | Tiff |
| | *Image bit depth* | 8-bit integer |
| | *When to save* | Every cycle |
| **ExportToSpreadsheet** | *Select the column delimiter* | Comma(",") |
| | *Add image metadata to your object data file?* | Yes |
| | *Press button to select measurements* | Nuc<br>- AreaShape/Refer to **features** for PCA<br>- Intensity/MinIntensity<br>- Intensity/MedianIntensity<br>- Number/Object<br>Cell<br>- AreaShape/Refer to **features** for PCA |

### High-content analysis of morphological features and GLB1/Bmi1 expression

| Module | Parameter | Setting |
|---|---|---|
| **Metadata** | *Refer to Proliferation of cAD-MSCs pipeline* - **Metadata** | |
| **NamesAndTypes** | *Select the rule criteria* | File Does Contain "DAPI" |
| | *Name to assign these images* | DAPI |
| | *Select the rule criteria* | File Does Contain "FITC" |
| | *Name to assign these images* | GLB1 |
| | *Select the rule criteria* | File Does Contain "Cy3" |
| | *Name to assign these images* | Skel |
| | *Select the rule criteria* | File Does Contain "Cy5" |
| | *Name to assign these images* | Bmi1 |
| | *Select the image type* | Grayscale image |
| | *Select intensity range from* | Image metadata |
| **Groups** | *Refer to Proliferation of cAD-MSCs pipeline* -**Groups** | |
| **IdentifyPrimaryObjects** | *Refer to HCA of morphological features and Ki67/Bmi1 -* **IdentifyPrimaryObjects** | |
| **IdentifySecondaryObjects** | *Refer to HCA of morphological features and Ki67/Bmi1 -* **IdentifySecondaryObjects** | |
| **EnhanceOrSuppressFeatures** | *Select the input image* | GLB1 |
| | *Name the output image* | EnhanceGLB1 |
| | *Select the operation* | Enhance |
| | *Feature type* | Speckles |
| | *Feature size* | 20 |
| | *Speed and accuracy* | Slow |
| **MaskImage** | *Select the input image* | EnhanceGLB1 |
| | *Name the output image* | MaskGLB1 |
| | *Use objects or an image as a mask* | Objects |
| | *Select object formask* | Cell |
| | *Invert the mask?* | No |
| **IdentifyPrimaryObjects** | *Use advanced settings* | Yes |
| | *Select the input image* | MaskGLB1 |
| | *Primary objects to be identified* | GLB1_Foci |
| | *Typical diameter of objects (Min; Max)* | 1;20 |
| | *Discard objects outside the diameter range?* | Yes |
| | *Discard objects touching the border of the image?* | No |
| | *Threshold strategy* | Adaptive |
| | *Thresholding method* | Otsu |
| | *Two-class or three-class thresholding* | Two classes |
| | *Threshold smoothing scale* | 0 |
| | *Threshold correction factor* | 1 |
| | *Lower and upper bounds on threshold* | 0.008;1.0 |
| | *Method to distinguish clumped objects* | Intensity |
| | *Method to draw dividing lines beween clumped objects* | Intensity |
| | *Automatically calculate size of smoothing filter for declumping* | No |
| | *Size of smoothing filter* | 0 |
| | *Automatically calculate minimum allowed distance between local maxima* | No |
| | *Suppress local maxima that are closer than this minimum allowed distance* | 2 |
| | *Speed up by using lower-resolution image to find local maxima* | No |
| | *Fill holes in identified objects?* | After both thresholding and declumping |

| | | |
|---|---|---|
| **RelateObjects** | *Parent objects* | Cell |
| | *Child objects* | GLB1_Foci |
| | *Name the output object* | GLB1_Foci_Cell |
| **MeasureObjectSizeShape** | *Select objects to measure* | Nuc, Cell |
| **MeasureObjectIntensity** | *Select an image to measure* | DAPI, Bmi1 |
| | *Select objects to measure* | Nuc, Cell |
| **RescaleIntensity** | *Select the input image* | DAPI |
| | *Name the output image* | ResDNA |
| | *Rescaling method* | Stretch each image to use the full intensity range |
| **RescaleIntensity** | *Select the input image* | Skel |
| | *Name the output image* | ResSkel |
| | *Rescaling method* | Stretch each image to use the full intensity range |
| **RescaleIntensity** | *Select input image* | GLB1 |
| | *Name the output image* | ResGLB1 |
| | *Rescaling method* | Stretch each image to use the full intensity range |
| **GrayToColor** | *Select a color scheme* | RGB |
| | *Select the image to be colored red* | ResSkel |
| | *Select the image to be colored green* | |
| | *Select the image to be colored blue* | ResDAPI |
| | *Name of the output image* | Fused |
| **OverlayOutlines** | *Select image on which to display outlines* | Fused |
| | *Name the output image* | SkelOver |
| | *Outline display mode* | Color |
| | *How to outline* | Outer |
| | *Select objects to display* | Nuc, Cell, GLB1_Foci_Cell |
| **SaveImages** | *Select the type of image to save* | Image |
| | *Select the image to save* | SkelOver |
| | *Select method for constructing file names* | From image filename |
| | *Saved file format* | Tiff |
| | *Image bit depth* | 8-bit integer |
| | *When to save* | Every cycle |
| **ExportToSpreadsheet** | *Select the column delimiter* | Comma(",") |
| | *Add image metadata to your object data file?* | Yes |
| | *Press button to select measurements* | Nuc<br>- AreaShape/Refer to **features** for PCA<br>- Intensity/MinIntensity<br>- Intensity/MedianIntensity<br>- Number/Object<br>Cell<br>- AreaShape/Refer to **features** for PCA<br>- Children/GLB1/Foci/Count |

### NEON transfection efficiency

| Module | Parameter | Setting |
|---|---|---|
| **Metadata** | *Refer to Proliferation of cAD-MSCs pipeline* - **Metadata** | |
| **NamesAndTypes** | *Select the rule criteria* | File Does Contain "Brightfield" |
| | *Name to assign these images* | BF |
| | *Select the rule criteria* | File Does Contain "FITC" |
| | *Name to assign these images* | NEON |
| | *Select the image type* | Grayscale image |
| | *Select intensity range from* | Image metadata |
| **Groups** | *Refer to Proliferation of cAD-MSCs pipeline* -**Groups** | |
| **RescaleIntensity** | *Select the input image* | BF |
| | *Name the output image* | RescaleBF |
| | *Rescaling method* | Stretch each image to use the full intensity range |
| **CorrectIlluminationCalculate** | *Select the input image* | RescaleBF |
| | *Name the output image* | IllumBF |
| | *Select how the illuiation fucntion is calculated* | Regular |
| | *Dilate objects in the final averaged image* | No |
| | *Rescale the illumination function* | Yes |
| | *Calculate function for each image individually?* | Each |
| | *Smoothing method* | Gaussian Filter |
| | *Method to calculate smoothing filter size* | Manually |
| | *Smoothing filter size* | 100 |
| **CorrectIlluminationApply** | *Select the input image* | RescaleBF |
| | *Name the output image* | CorrBF |
| | *Select the illumination funciton* | IllumBF |
| | *Select how the function is applied* | Divide |
| **ImageMath** | *Operation* | Invert |
| | *Name the output image* | MathBF |
| | *Select the first image* | CorrBF |
| | *Multiply the first image by* | 1.0 |
| | *Raise the power of the result by* | 1.0 |
| | *Multiply the result by* | 1.0 |
| | *Add to result* | 0.0 |
| | *Set values less than 0 equal to 0* | Yes |
| | *Set values greater than 1 equal to 1* | Yes |
| | *Ignore the image masks* | Yes |
| **EnhanceOrSuppressFreatures** | *Select the input image* | MathBF |
| | *Name the output image* | EnhanceBF |
| | *Select the operation* | Enhance |
| | *Feature type* | Speckles |
| | *Feature size* | 50 |
| | *Speed and accuracy* | Fast |
| **IdentifyPrimaryObjects** | *Use advanced settings* | Yes |
| | *Select the input image* | EnhanceBF |
| | *Primary objects to be identified* | CellBF |
| | *Typical diameter of objects (Min; Max)* | 35;100 |
| | *Discard objects outside the diameter range?* | Yes |
| | *Discard objects touching the border of the image?* | No |
| | *Threshold strategy* | Global |
| | *Thresholding method* | Minimum cross entropy |
| | *Threshold smoothing scale* | 1.3488 |
| | *Threshold correction factor* | 1 |
| | *Lower and upper bounds on threshold* | 0.08;1.0 |

| | | |
|---|---|---|
| | *Method to distinguish clumped objects* | Intensity |
| | *Method to draw dividing lines beween clumped objects* | Propagate |
| | *Automatically calculate size of smoothing filter for declumping* | Yes |
| | *Automatically calculate minimum allowed distance between local maxima* | No |
| | *Suppress local maxima that are closer than this minimum allowed distance* | 10 |
| | *Speed up by using lower-resolution image to find local maxima* | Yes |
| | *Fill holes in identified objects?* | After both thresholding and declumping |
| **MeasureObjectSizeShape** | *Select objects to measure* | CellBF |
| **MeasureObjectIntensity** | *Select an image to measure* | NEON |
| **OverlayOutlines** | *Select image on which to display outlines* | CorrBF |
| | *Name the output image* | BFOver |
| | *Outline display mode* | Color |
| | *How to outline* | Thick |
| | *Select objects to display* | CellBF |
| **SaveImages** | *Select the type of image to save* | Image |
| | *Select the image to save* | BFOver |
| | *Select method for constructing file names* | From image filename |
| | *Saved file format* | Tiff |
| | *Image bit depth* | 8-bit integer |
| | *When to save* | Every cycle |
| **ExportToSpreadsheet** | *Select the column delimiter* | Comma(",") |
| | *Add image metadata to your object data file?* | Yes |
| | *Press button to select measurements* | CellBF<br>- Intensity/MedianIntensity<br>- Intensity/MinIntensity |

### Nuclear and cellular features for principal component analysis

| Nuc/Cell features[3] |
|---|
| Area |
| Compactness |
| Eccentricity |
| Form factor |
| Major axis length |
| Minor axis length |
| Maximal Feret diameter |
| Median radius |
| Perimeter |
| Solidity |

[3] For detailed feature definition: http://cellprofiler-manual.s3.amazonaws.com/CellProfiler-3.1.8/index.html
