## Supplementary Table S3 for "High-content analysis reveals senescence of cultured canine adipose-derived mesenchymal stromal cells to be reversible with a novel immortalization approach"

### Supplementary Table S3: PCR and immunostaining protocols

#### *PCR primers*

| PROCEDURE | NAME | FW | REV |
|---|---|---|---|
| **Infusion Cloning** | hBmi1-hTERT | CGGCCCTATGCATCGAACAACGAGAATCAAG | TCCAAGCTTATGTACATTATCAGCCAGGATGGTCTTGAAG |
| | Neon | ATCACACTGGCGGCCGCCATGGTGAGCAAGGGCGAGGAG | CGATGCATAGGGCCGGGATTCTCTTCCAC |
| | Screen | CAAAGGAAAAGGGCCTTTC | CGGAGCCATCTACCATGG |
| **Immortalization** | Neon | TGGCTTCCATCAGTACCTGC | AGTCTTCTTCGACCTGCACC |
| | Transition | GAGTGACAAGGCCAACAGCC | CAGCACCTCGCGGTAGT |
| | hTERT | GCAGCTACCTGCCCAACACG | CACACCTGGTAGGCGCAG |
| | cBmi1 | ACCCTATGTTTGAGCCTTATCAG | CAAACTGAAAACGATTATGGTTTTTCG |
| | GAPDH | GCCTCCTGCACCACCAAC | CATACCAGGAAATGAGCTTG |

#### *PCR protocol*

| PROCEDURE | REAGENT/METHOD | TIME/TEMPERATURE |
|---|---|---|
| **Cell harvesting** | TRIzol® RNA Isolation Reagent | ice-cold |
| **Cell storage** | TRIzol® RNA Isolation Reagent | -80°C |
| **Sample thawing** | TRIzol® RNA Isolation Reagent | RT |
| **Sample homogenisation** | 30G syringe needle | RT |
| **RNA extraction** | Chloroform (Merck), 20% of the sample volume, centrifugation 12'000 x *g* | 15 min, 4°C |
| **Aqueous upper phase collection** | Collection in clean tubes and addition of the same volume of 70% ethanol (Merck) | RT |
| **RNA purification** | PureLink® RNA Mini Kit | RT |
| **DNAse** | PureLink® DNase Set | RT |
| **Reverse transcription** | iScript™ cDNA Synthesis Kit | RT |
| **PCR** | Platinum™ Taq Green Hot Start Ploymerase Kit | Annealing T=60°C |
| **Gel electrophoresis** | 2% UltraPure agarose gels, SYBR® Safe DNA Gel Stain | RT |

*Unless stated otherwise, the products in this table were purchased from ThermoFisher Scientific, Reinach, CH*

***Immunostaining antibodies***

| 1st Ab | 2nd Ab | Imaging filter set |
|---|---|---|
| 1:400 mouse anti-Ki67(20Raj1)[1] | 1:200 goat anti-mouse Alexa Fluor®488[2] | 490_20x & 525_36m |
| 1:100 mouse anti-GLB1 (OTI10B2)[3] | 1:200 goat anti-mouse Alexa Fluor®488[2] | 490_20x & 525_36m |
| 1:200 rabbit anti-Bmi1 (AA276-326)[4] | 1:200 goat anti-rabbit Alexa Fluor®647[2] | 645_30x & 705_72m |

1 µg/ml DAPI[5]: imaging filter set: 350_50x & 455_50m
ActinRed™ 555 Ready Probes™ Reagent (phalloidin)[6]: imaging filter set 543_22x & 605_64m

***Immunostaining protocol***

| Procedure | Reagent | Time |
|---|---|---|
| Wash 2x | PBS | 2x5min |
| Fixation | 2% paraformaldehyde (PFA/PBS) | 20 min |
| Wash 2x | PBS | 2x5 min |
| Permeabilization | 0.1% Triton X-100[7] | 15 min |
| Blocking | Dako Serum-free Protein Block solution[8] | 30 min |
| Primary antibody | Diluted in 0.05% TWEEN® 20 in PBS | Overnight (at 4°C) |
| Wash 2x | PBS | 2x5min |
| Secondary antibody | Diluted in 0.05% TWEEN® 20 in PBS | 2 h |
| Wash 2x | PBS | 2x5min |
| Nuclei and cell contour staining | 1µg/ml DAPI[5], Alexa Fluor™ 555 phalloidin[6] | 20 min |
| Wash 2x | PBS | 2x5min |

*Unless stated otherwise, the procedures were performed at RT*

[1] #14-5699-95, eBioscience™ThermoFisher Scientific, Reinach, CH
[2] #115-546-003 and #111-605-144, Jackson ImmunoResearch Laboratories Inc., West Grove, PA, USA
[3] # TA505625, OriGene, ThermoFisher Scientific, Reinach, CH
[4] ab85688, Abcam, Cambridge, UK
[5] D1306, Invitrogen™, ThermoFisher Scientific, Reinach, CH
[6] R37112, Invitrogen™, ThermoFisher Scientific, Reinach, CH
[7] 108643, Merck, Zug, CH
[8] X090930-2 , Agilent, Santa Clara, CA, USA
