## Supplementary Table S4 for "High-content analysis reveals senescence of cultured canine adipose-derived mesenchymal stromal cells to be reversible with a novel immortalization approach"

### Supplementary Table S4: Numerical results

| Output | Unit | Cells | P2 | P3 | P6 | P9 |
|---|---|---|---|---|---|---|
| **3.1 Proliferation** | | | | | | |
| Nuclei count DIV7 | x10$^3$ | cAD-MSCs | 81.7 ±42.6<br>n = 15 | 102.2 ±7.6<br>n = 15 | **32.9** ±12.6<br>n = 15<br>p = 1.5e-6 | **3.0** ±2.4<br>n = 15<br>p = 1.3 e-8 |
| | | Hela | 103.4 ±7.2<br>n = 3 | 82.4 ±3.8<br>n = 3 | 54.8 ±5.8<br>n = 3 | 71.1 ±14.9<br>n = 3 |
| | | HelaSen | 0.8 ±0.2<br>n = 3 | 2.0 ±0.7<br>n = 3 | 0.3 ±0.2<br>n = 3 | 5.2 ±1.6<br>n = 3 |
| eAUC | x10$^3$ | cAD-MSCs | 79.9 ±38.4<br>n = 5 | **186.0**±67.2<br>n = 5<br>p = 0.016 | **35.2** ±24.8<br>n = 5<br>p = 7.9e-3 | **2.5** ±2.9<br>n = 5<br>p = 7.9e-3 |
| *K* | x10$^3$ | cAD-MSCs | 60.1 ±38.5<br>n = 4 | 116.0 ±11.8<br>n = 5 | **21.4** ±13.3<br>n = 5<br>p = 7.9e-3 | **3.19** ±2.58<br>n = 4<br>p = 0.016 |
| **3.2 Morphology** | | | | | | |
| Nuclear area | 10$^2$µm$^2$ | cAD-MSCs | 0.69 ±0.21<br>n=773 | **0.76** ±0.26<br>n=1381<br>p ~ 0 | **0.86** ±0.22<br>n=700<br>p ~ 0 | **1.31** ±0.60<br>n=68<br>p ~ 0 |
| Cellular area | 10$^2$µm$^2$ | cAD-MSCs | 3.11 ±1.44<br>n=773 | **2.95** ±1.56<br>n=1381<br>p =3.7e-8 | **5.45** ±3.56<br>n=700<br>p ~ 0 | **28.6** ±22.2<br>n=68<br>p =2.7e-13 |
| **3.3 Protein expression** | | | | | | |
| Median Ki67 intensity | AIU x10$^{-3}$ | cAD-MSCs | 1.7 ±0.4<br>n=312 | **2.7** ±1.6<br>n=550<br>p ~ 0 | **1.5** ±0.5<br>n=322<br>p ~ 0 | 1.4 ±0.6<br>n=30 |
| Median Bmi1 intensity | AIU x10$^{-3}$ | cAD-MSCs | 5.2 ±1.4<br>n=710 | **8.8** ±2.8<br>n=1214<br>p ~ 0 | **6.0** ±1.5<br>n=615<br>p ~ 0 | **7.1** ±2.3<br>n=54<br>p = 7.2e-5 |
| Bmi1 positive cells | % | cAD-MSCs | 37.2 ±13.0<br>n=5 | **100** ±1.7<br>n=5<br>p = 7.9e-3 | **70.6** ±6.9<br>n=5<br>p = 7.9e-3 | **85.7** ±5.9<br>n=5<br>p = 0.016 |
| Ki67 positive cells | % | cAD-MSCs | 57.9 ±5.7<br>n=5 | **29.8** ±10.8<br>n=5<br>p = 7.9e-3 | 22.1 ±2.9<br>n=5 | **52.0** ±26.4<br>n=5<br>p = 0.032 |
| **3.4 Immortalization** | | | | | | |
| Nuclei count DIV7 | x10$^3$ | Immo Dog | NA | NA | NA | 18.0 ±1.0<br>n=3 |
| | | GFP Dog | NA | NA | NA | 5.5 ±1.0<br>n=3 |
| **3.5 Immortalization morphology and protein expression** | | | | | | |
| Nuclear area | 10$^2$µm$^2$ | Immo Dog | NA | NA | 2.3 ±0.5<br>n=5838 | **2.2** ±0.5<br>n=3422<br>p = 5.3e-7 |
| Cellular area | 10$^2$µm$^2$ | Immo Dog | NA | NA | 64.1 ±34.0<br>n=5838 | **55.4** ±27.2<br>n=3422<br>p ~ 0 |
| Median Ki67 intensity | AIU x10$^{-3}$ | Immo Dog | NA | NA | 1.9 ±0.4<br>n=2027 | **7.8** ±3.0<br>n=1136<br>p = 3.7e-5 |
| Median Bmi1 intensity | AIU x10$^{-3}$ | Immo Dog | NA | NA | 20.0 ±22.0<br>n=1881 | **63.0** ±52.0<br>n=1200<br>p ~ 0 |

All the presented data are mean values with standard deviation in grey. The number of samples (n) is reported for each measurement. Bold characters represent values that significantly differ ($p < 0.05$) from the precedent measured passage under the same condition. Only $p<0.05$ are reported in detail in red.
NA=not available; this time point was not measured.

**Statistical procedure**

***3.1 Proliferation***

- cAD-MSCs: 5 biological replicates (Dog1-5) and 3 technical replicates for each biological replicate and passage P
- Hela and HelaSen: 3 technical replicates per each passage P

Per each cell line, the different passages where compared using the non-parametric Kruskal-Wallis multiple comparison test with a Wilcoxon post hoc test using Bonferroni correction.

***3.2 Morphology***

- cAD-MSCs: 5 biological replicates (Dog1-5) and 12 technical replicates for each biological replicate and passage P

After observing a problem with overpower which resulted in significant differences between the biological samples, we performed stratified subsampling. After subsampling no major difference between the biological replicates was observed within a passage (Supplementary Figure S1), therefore the data were pooled for further analysis. Notably, after subsampling, the differences between the passages were maintained.
The different passages were compared using the non-parametric Kruskal-Wallis multiple comparison test with a Wilcoxon post hoc test using Bonferroni correction. This was due to the observation that a normal distribution could not be assumed (Shapiro-Wilk test) due to a strong skewness of the data.

***3.3 Protein expression***

- cAD-MSCs: 5 biological replicates (Dog1-5) and 4 technical replicates for each biological replicate and passage P for Ki67 and 8 technical replicates for Bmi1

For the evaluation of the median intensity values per cell, the biological replicates were pooled, while for the percentage of the fluorescence positive cells, the mean of the replicates per each biological samples is further analyzed and plotted per passage.

***3.4 Immortalization***

- Immo Dog and GFP Dog: for this experiment, a single dog sample was transfected with either one or the other condition. For all the experiments, 3 technical replicates were performed.

***3.5 Immortalization and protein expression***

- Immo Dog: for this experiment, a single dog sample was transfected with the immortalization plasmid. For the experiments related to morphology, 12 technical replicates were performed. For Ki67 intensity measurements, 4 technical replicates were included, while for Bmi1, 8 technical replicates were performed.

The two different passages were compared using the non-parametric Wilcoxon rank-sum test as normal distribution was not given due to a moderate skewness of the data (Shapiro-Wilk test).

Post hoc power analysis using the G*Power 3.1.9.7 software revealed that for each experiment a power $> 0.95$ was reached.
